# Real time monitoring of peptidoglycan synthesis by membrane-reconstituted penicillin binding proteins

**DOI:** 10.1101/2020.08.02.233189

**Authors:** Víctor M. Hernández-Rocamora, Natalia Baranova, Katharina Peters, Eefjan Breukink, Martin Loose, Waldemar Vollmer

**Affiliations:** Centre for Bacterial Cell Biology, Biosciences Institute, Newcastle University, Richardson Road, Newcastle upon Tyne, NE2 4AX, UK; Institute for Science and Technology Austria (IST Austria), Klosterneuburg, Austria; Membrane Biochemistry and Biophysics, Bijvoet Centre for Biomolecular Research, University of Utrecht, Padualaan 8, 3584 Utrecht, The Netherlands

**Author notes:** Contributed equally (Víctor M. Hernández-Rocamora, Natalia Baranova).

## Abstract

Peptidoglycan is an essential component of the bacterial cell envelope that surrounds the cytoplasmic membrane to protect the cell from osmotic lysis. Important antibiotics such as β-lactams and glycopeptides target peptidoglycan biosynthesis. Class A penicillin binding proteins are bifunctional membrane-bound peptidoglycan synthases that polymerize glycan chains and connect adjacent stem peptides by transpeptidation. How these enzymes work in their physiological membrane environment is poorly understood. Here we developed a novel FRET-based assay to follow in real time both reactions of class A PBPs reconstituted in liposomes or supported lipid bilayers and we demonstrate this assay with PBP1B homologues from *Escherichia coli, Pseudomonas aeruginosa* and *Acinetobacter baumannii* in the presence or absence of their cognate lipoprotein activator. Our assay allows unravelling the mechanisms of peptidoglycan synthesis in a lipid-bilayer environment and can be further developed to be used for high throughput screening for new antimicrobials.

## INTRODUCTION

Peptidoglycan (PG) is a major cell wall polymer in bacteria. It is composed of glycan strands of alternating N-actetylglucosamine (Glc*N*Ac) and N-acetylmuramic acid (Mur*N*Ac) residues interconnected by short peptides. PG forms a continuous, mesh-like layer around the cell membrane to protect the cell from bursting due to the turgor and to maintain cell shape (Vollmer *et al*., 2008). The essentiality and conservation of PG in bacteria make peptidoglycan metabolism an ideal target of antibiotics.

Class A penicillin-binding proteins (PBPs) are bifunctional PG synthases, which uses the precursor lipid II to polymerize glycan chains (glycosyltransferase reactions) and crosslink peptides from adjacent chains by DD-transpeptidation (Goffin & Ghuysen, 1998). Moenomycin inhibits the glycosyltransferase and β-lactams the transpeptidase function of class A PBPs (Sauvage & Terrak, 2016, Macheboeuf *et al*., 2006). In *E. coli*, PBP1A and PBP1B account for a substantial proportion of the total cellular PG synthesis activity (Cho *et al*., 2016) and they are tightly regulated by interactions with multiple proteins (Egan *et al*., 2015, Typas *et al*., 2012, Egan *et al*., 2017, Egan *et al*., 2020), including the outer membrane anchored activators LpoA and LpoB (Egan *et al*., 2018, Typas *et al*., 2010, Jean *et al*., 2014).

Historically, *in vitro* PG synthesis assays have been crucial to decipher the biochemical reactions involved in PG synthesis and determine the mode of action of antibiotics (Izaki *et al*., 1968). However, these studies were limited by the scarcity of lipid II substrate and the inability to purify a sufficient quantity of active enzymes. Lipid II can now be synthesized chemically (VanNieuwenhze *et al*., 2002, Schwartz *et al*., 2001, Ye *et al*., 2001) or semi-enzymatically (Breukink *et al*., 2003, Egan *et al*., 2015), or isolated form cells with inactivated MurJ (Qiao *et al*., 2017). Radioactive or fluorescent versions of lipid II are also available to study PG synthesis in the test tube. However, there are several drawbacks with currently available PG synthesis assays. First, most assays are end-point assays that rely on discrete sampling and therefore do not provide real-time information about the enzymatic reaction. Second, some assays involve measuring the consumption of lipid II or analysing the reaction products by SDS-PAGE (Egan *et al*., 2015, Barrett *et al*., 2007, Qiao *et al*., 2014, Sjodt *et al*., 2018) or HPLC after digestion with a muramidase (Bertsche *et al*., 2005, Born *et al*., 2006). These laborious techniques make assays incompatible with high through-put screening and hinder the determination of kinetic parameters. A simple, real-time assay with dansyl-labelled lipid II substrate overcomes these problems but is limited to assay GTase reactions (Schwartz *et al*., 2001, Offant *et al*., 2010, Egan *et al*., 2015).

Recently two types of real-time TPase assays have been described. The first uses non-natural mimics of TPase substrates such as the rotor-fluorogenic 470 D-lysine probe Rf470DL, which increases its fluorescence emission upon incorporation into PG (Hsu *et al*., 2019). The second assay monitors the release of D-Ala during transpeptidation in coupled enzymatic reactions with D-amino acid oxidase, peroxidases and chromogenic or fluorogenic compounds (Frere *et al*., 1976, Gutheil *et al*., 2000, Catherwood *et al*., 2020). Coupled assays are often limited in the choice of the reaction conditions, which in this case must be compatible with D-amino acid oxidase activity. Hence, each of the current assays has its limitations and most assays exclusively report on either the GTase or TPase activity, but not both activities at the same time.

Another major drawback of many of the current assays is that they include detergents and/or high concentration (up to 30%) of the organic solvent dimethyl sulfoxide (DMSO) to maintain the PG synthases in solution (Offant *et al*., 2010, Biboy *et al*., 2013, Huang *et al*., 2013, Lebar *et al*., 2013, Qiao *et al*., 2014, Egan *et al*., 2015, Catherwood *et al*., 2020). However, both detergents and DMSO have been shown to affect the activity and interactions of *E. coli* PBP1B (Egan & Vollmer, 2016). Importantly, a freely diffusing, detergent-solubilised membrane enzyme has a very different environment compared to the situation in the cell membrane where it contacts phospholipids and is confined in two dimensions (Gavutis *et al*., 2006, Zhdanov & Höök, 2015). Here we sought to overcome the main limitations of current PG synthesis assays. We established the sensitive Förster Resonance Energy Transfer (FRET) detection technique for simultaneous monitoring of GTase and TPase reactions. The real-time assay reports on PG synthesis in phospholipid vesicles or planar lipid bilayers. We successfully applied this assay to several class A PBPs from pathogenic Gram-negative bacteria, demonstrating its robustness and potential use in screening assays to identify PBP inhibitors.

## RESULTS

### Real time assay for detergent-solubilised *E. coli* PBP1B

To develop a FRET-based real time assay for PG synthesis using fluorescently labelled lipid II, we prepared lysine-type lipid II versions with high quantum yield probes, Atto550 (as FRET donor) and Atto647n (as FRET acceptor), linked to position 3 (Figure 1 – figure supplement 1A-B) (Mohammadi *et al*., 2014, Egan *et al*., 2015). For assay development we used *E. coli* PBP1B (PBP1B^Ec^) (Egan *et al*., 2015, Bertsche *et al*., 2005, Biboy *et al*., 2013) solubilized with Triton X-100 and a lipid-free version of its cognate outer membrane-anchored lipoprotein activator LpoB (Typas *et al*., 2010, Egan *et al*., 2014, Egan *et al*., 2018, Lupoli *et al*., 2014, Catherwood *et al*., 2020).

PBP1B^Ec^ can utilize fluorescently labelled lipid II to polymerize long glycan chains only when unlabelled lipid II is also present in the reaction (van&t Veer, 2016). We therefore included unlabelled *m*DAP-type lipid II into reactions of PBP1B^Ec^ with lipid II-Atto550 and lipid II-Atto647n, with or without LpoB(sol) (Figure 1A). We monitored reactions by measuring fluorescence intensities in real time in a microplate reader for 60 min (Figure 1B, Figure 1 – figure supplement 1C). Consistent with fluorescence transfer occurring, the emission of the acceptor fluorophore at 665 nm (FI_acceptor_) increased while the emission of the donor fluorophore at 580 nm (FI_donor_) decreased, giving rise to an increase in the FI_acceptor_/FI_donor_ ratio (Figure 1 – figure supplement 1C). No FRET was observed in samples containing the GTase inhibitor moenomycin, which indirectly also inhibits TPase reactions (Bertsche *et al*., 2005) (Figure 1 – figure supplement 1C). Without LpoB, FRET appeared after ∼5 min and slowly increased until it plateaued after 50-60 min (Figure 1B). By contrast, reactions with LpoB(sol) showed an immediate and rapid increase in FRET which reached the plateau after 10-20 minutes, consistent with faster PG synthesis (Figure 1B, left panel). The presence of the TPase inhibitor ampicillin generally reduced the final FRET level by ∼3-fold (Figure 1B, middle panel), indicating that the FRET is mainly a result of TPase reactions. As expected, ampicillin did not prevent the stimulation of PBP1B^Ec^ by LpoB(sol) which accelerated the FRET increase by 10-20 times with or without ampicillin (Figure 1C), consistent with the previously reported stimulation of both, GTase and TPase activities (Typas *et al*., 2010, Egan *et al*., 2014, Egan *et al*., 2018).

**Figure 1.**
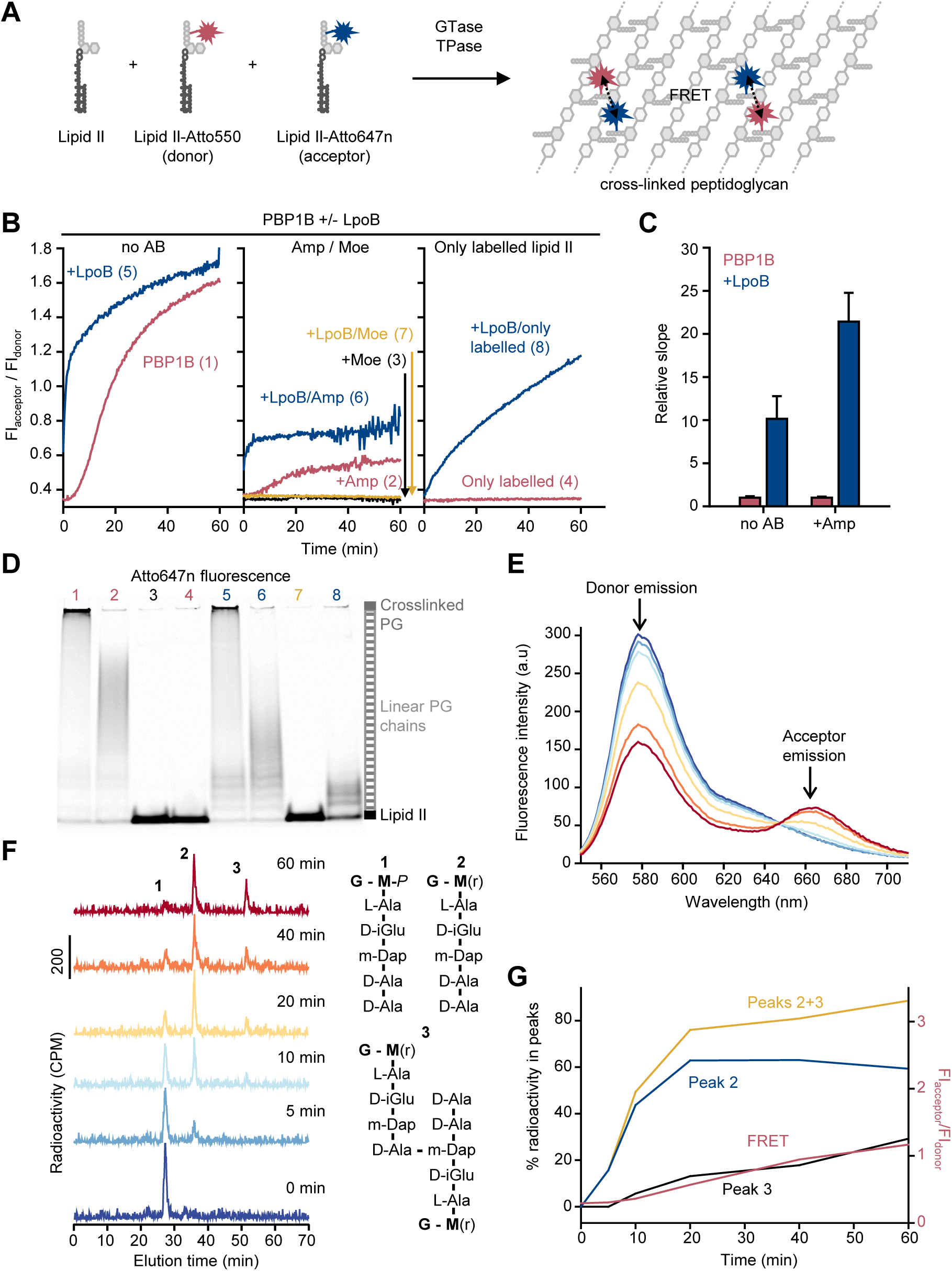
FRET assay to monitor peptidoglycan synthesis in real time. (**A**) Scheme of the reactions of a class A PBP (GTase-TPase) with unlabelled lipid II and the two versions of labelled lipid II, yielding a PG product that shows FRET. (**B**) Representative reactions curves from FRET assays of detergent-solubilised PBP1B^Ec^. The enzyme (0.5 µM) was mixed with unlabelled lipid II, Atto550-labelled lipid II and Atto647n-labelled lipid II at a 1:1:1 molar ratio (each 5 µM), in the presence or absence of 2 µM LpoB(sol). Reactions were performed in the absence of antibiotic (left panel), with 1 mM ampicillin (Amp) or 50 µM moenomycin (Moe) (middle panel), or by omitting unlabelled lipid II (right panel). The numbers indicate the corresponding lane of the gel in panel **D**. Samples were incubated for 1 h at 25°C. (**C**) Averaged initial slopes from reaction curves obtained by the FRET assay for detergent-solubilised *E. coli* PBP1B in the presence (blue) or absence (red) of LpoB, and in the presence or absence of ampicillin. Values are normalised relative to the slope in the absence of activator for each condition and are mean ± SD of 2-3 independent experiments. (**D**) Aliquots at the end of the reactions shown in **B** were boiled and analysed by SDS-PAGE using fluorescence detection of the acceptor (Atto647n), lanes are labelled with the reaction numbers in **B**. (**E**) and (**F**), PBP1B^Ec^ (0.5 µM) was incubated with 5 µM each of lipid II-Atto647n, lipid II-Atto550 and ^14^C-labelled lipid II. At indicated time points, aliquots were taken and reactions were stopped by addition of moenomycin. After measuring fluorescence (**E**), the PG was digested with the muramidase cellosyl, and the resulting muropeptides were reduced with sodium borohydride and separated by HPLC (**F**). The structures of muropeptides corresponding to peaks 1-3 are shown next to the chromatograms. (**G**) Quantification of peak 2 (GTase product, blue), peak 3 (GTase+TPase, black) or the sum of both 2 and 3 (yellow) from chromatograms in **F**, along with the FRET signal (red) calculated as ratio of acceptor emission over donor emission from data in **E**.

We also analysed the reaction products by SDS-PAGE combined with fluorescence scanning. This analysis confirmed the formation of PG chains containing both fluorophores, Atto550 and Atto647n and that ampicillin blocked the formation of cross-linked PG and moenomycin inhibited glycan strand formation (Figure 1D, Figure 1 – figure supplement 1D).

### Intra-chain versus inter-chain FRET

Because ampicillin substantially reduced the FRET signal we hypothesized that FRET arises mainly between probe molecules residing on different glycan chains of a cross-linked PG product (Figure 1 – figure supplement 1A). To determine the contribution of intra-chain FRET, we performed reactions with only labelled lipid II (Figure 1B, right panel), where cross-linking is not possible. Without LpoB(sol), PBP1B^Ec^ was unable to use lipid II-Atto550 and lipid II-Atto647n for polymerization (Figure 1B, D), confirming a previous study (van&t Veer, 2016). In the presence of LpoB(sol), PBP1B^Ec^ produced short, non-crosslinked individual PG chains (Figure 1D) that gave rise to a slow but large increase in FRET (Figure 1B, right panel; Figure 1 – figure supplement 1C), indicating that lipid II polymerization reactions occurred. Our combined data also suggest that the total FRET signal emerges from two steps that have different rates. First, the formation of linear glycan chain causes initially a slow and moderate FRET increase and second, once peptide cross-linking reactions begin, the FRET increases fast and reaches a high level.

To confirm that the formation of peptide crosslinks is required to produce substantial FRET in the absence of LpoB, we analysed the PG synthesised by PBP1B^Ec^ from radioactively labelled lipid II and the two fluorescent lipid II analogues (Figure 1E-G). We monitored the reaction at different time points by fluorescence spectroscopy (FRET measurements) and digested aliquots with the muramidase cellosyl before separating the resulting muropeptides by HPLC. The monomers and cross-linked muropeptide dimers were quantified by scintillation counting using an in-line radiation detector attached to the HPLC column (Figure 1F). FRET increased over time and correlated well with the formation of cross-linked muropeptide dimers, but not the rate of lipid II consumption (peak 2) (Figure 1G). Overall, we conclude that the FRET assay is capable of reporting GTase activity alone, but the overall FRET signal is dominated by the TPase activity.

### FRET assay to monitor PG synthesis in liposomes

To establish the FRET assay for membrane-embedded PG synthases we reconstituted a version of PBP1B^Ec^ with a single cysteine at the cytoplasmic N-terminus into liposomes prepared from *E. coli* polar lipids (EcPL) (Figure 2 – figure supplement 1A). The liposome-reconstituted PBP1B^Ec^ became accessible to a sulfhydryl-reactive fluorescent probe only after disrupting the liposomes with detergent (Figure 2A), showing that virtually all PBP1B molecules were oriented with the N-terminus inside the liposomes. Next, we reconstituted unmodified PBP1B^Ec^ and tested its activity by adding radioactive lipid II. In contrast to the detergent-solubilized enzyme, the liposome-reconstituted PBP1B^Ec^ required the absence of NaCl from the reaction buffer for improved the activity (Figure 2 – figure supplement 1B-E), suggesting that ionic strength affects either the structure of PBP1B^Ec^ in the membrane, the properties of EcPL liposomes or the delivery of lipid II into the liposomes.

**Figure 2.**
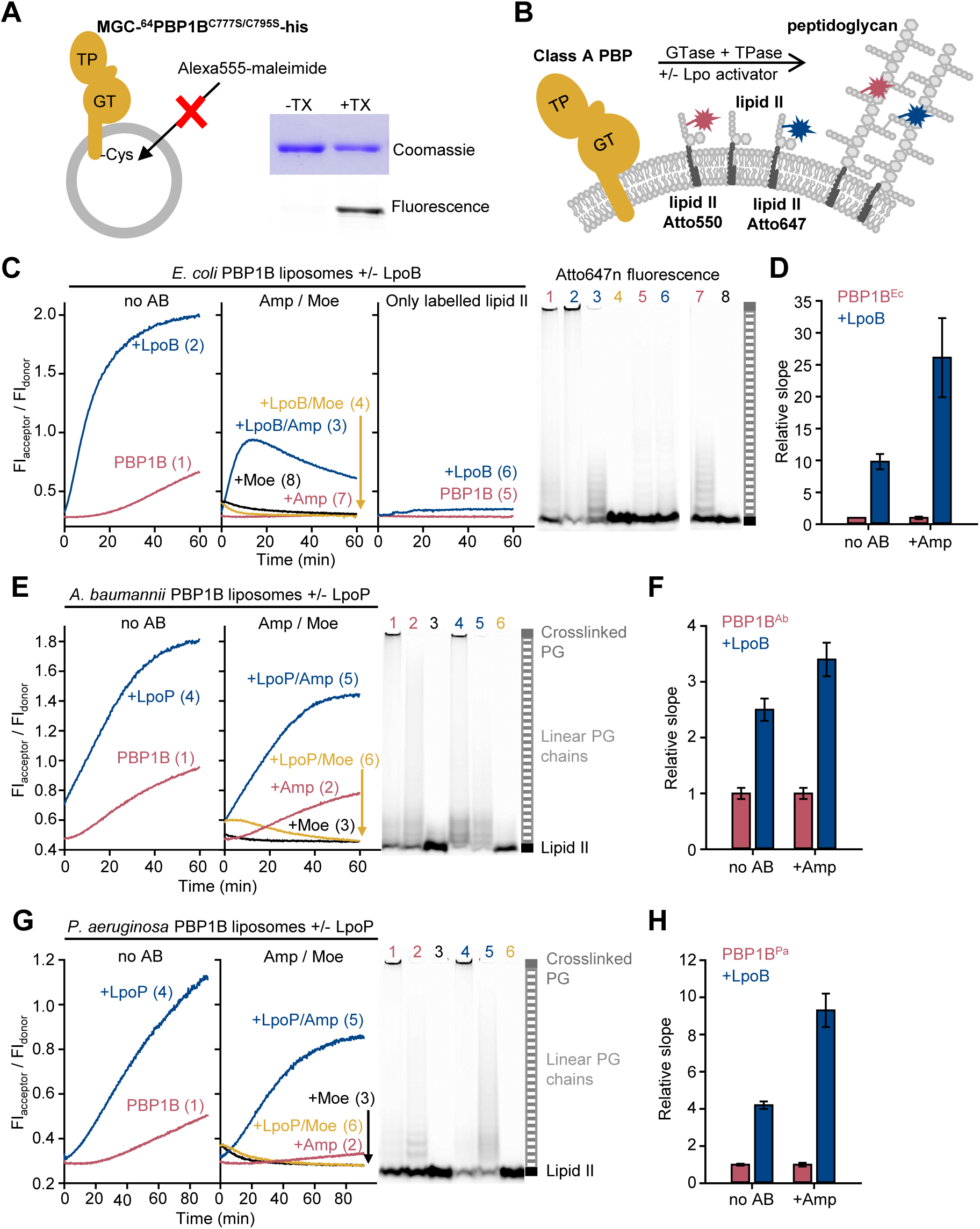
The FRET assay for PG synthesis can be adapted for reactions on liposomes. (**A**) Class A PBPs were reconstituted in *E. coli* polar lipid liposomes. To assess the orientation of the liposome-reconstituted PBPs, MGC-^64^PBP1B-his C777S C795S containing a single cysteine in the N-terminal region was reconstituted as in **A**. The accessibility of the cysteine was determined by staining with sulfhydryl-reactive fluorescent probe, AlexaFluor555-maleimide, in the presence or absence of Triton X-100 (TX). Samples were analysed by SDS-PAGE with fluorescence scanning to detect labelled protein followed by Coomassie staining. (**B**) To perform activity assays in liposomes, class A PBPs were reconstituted along a 1:1 molar ratio mixture of Atto550-labelled lipid II and Atto647n-labelled lipid II in liposomes as in **A**. Reactions were started by addition of unlabelled lipid II in the presence or absence of lipoprotein activators (lpo). Using this methodology, we monitored the activity of PBP1B^Ec^ (**C-D**), PBP1B^Ab^ (**E-F**) and PBP1B^Pa^ (**G**-**H**). Representative reactions curves are shown. Reactions were carried out in the presence (blue lines) or absence (red lines) of the lipoprotein activators (LpoB(sol) for PBP1B^Ec^, LpoP^Ab^(sol) for PBP1B^Ab^ and LpoP^Pa^(sol) for PBP1B^Pa^), and either in the absence of antibiotic (left) or in the presence of 1 mM ampicillin (Amp) or 50 µM moenomycin (Moe, black and yellow lines) (middle). For PBP1B^Ec^, control reactions in the absence of unlabelled lipid II (right panel) are also shown. Products were analysed by SDS-PAGE followed by fluorescence scanning at the end of reactions (right side). Curves are numbered according the corresponding lane on the SDS-PAGE gels. PBP1B^Ec^, PBP1B^Ab^ and PBP1B^Pa^ were reconstituted in EcPL liposomes containing labelled lipid II (0.5 mol% of lipids, 1:1 molar ratio mixture of atto550-labelled lipid II and Atto647n-labelled lipid II), at protein to lipid molar ratios of 1:3000, 1:2000 and 1:3000, respectively. Reactions were started by adding unlabelled lipid II (final concentration 12 µM) and incubated at 37°C for 60 min (PBP1B^Ec^ and PBP1B^Ab^) or 90 min (PBP1B^Pa^) while monitoring fluorescence at 590 and 680 nm with excitation at 522 nm. (**D**), (**F**) and (**H**) show averaged initial slopes from reaction curves obtained by the FRET assay for liposome-reconstituted PBP1B^Ec^, PBP1B^Ab^ and PBP1B^Pa^, respectively, in the presence (blue) or absence (red) of lipoprotein activators and in the presence or absence of ampicillin. Values are normalised relative to the slope in the absence of activator and are mean ± variation of 2 independent experiments.

We next aimed to adapt the FRET assay to study PG synthesis on liposomes (Figure 2, Figure 2 – figure supplement 2). As PBP1B^Ec^ did not accept Atto550-or Atto647-derivatised lipid II for GTase reactions in the absence of unlabelled lipid II (Figure 1B), we reconstituted PBP1B^Ec^ in liposomes along both Atto-labelled substrates and initiated the reaction by adding unlabelled lipid II (Figure 2B). PBP1B^Ec^ reaction rates in liposomes were slower than in the presence of Triton X-100 and we noticed a lag time before FRET started to increase (Figure 2C, left panel). Ampicillin or moenomycin blocked the increase in FRET (Figure 2C, middle panel). For an unknown reason, the FRET signal with moenomycin was initially higher than without moenomycin and then decreased to initial values without moenomycin (Figure 2C, middle panel), independent of the class A PBP used (see below) but not in empty liposomes (Figure 2 – figure supplement 3). LpoB(sol) produced a ∼10-fold increase in the initial slope (Figure 2D) and the resulting final FRET was higher (Figure 2C, left panel). Interestingly, in the presence of ampicillin and LpoB(sol), FRET increased rapidly at the start of reactions, but then decreased slowly, reaching a lower FRET value than in the presence of LpoB(sol) alone (without ampicillin) (Figure 2C, middle panel). The decrease in FRET in the presence of ampicillin suggests the spectroscopic properties of the incorporated probes change over time, presumably by moving them further away from the lipid end of the growing glycan chains. Liposomes without unlabelled lipid II produced a low FRET signal only in the presence of LpoB(sol) (Figure 2C, right panel). The analysis of the final products by SDS-PAGE confirmed that both Atto550 and Atto647n were incorporated into glycan chains or cross-linked peptidoglycan during the reaction in liposomes (Figure 2C, right side, Figure 2 – figure supplement 2B). In summary, we established a FRET-based assay that allows to monitor the activity of membrane-reconstituted PBP1B in real time and showed that the FRET signal was sensitive to the presence of PG synthesis inhibitors (moenomycin and ampicillin).

### Activities of other membrane-bound class A PBPs

To demonstrate the usefulness of the FRET assay to study class A PBPs of potential therapeutic interest, we next tested two PBP1B homologues from Gram-negative pathogens, *Acinetobacter baumannii* (PBP1B^Ab^) and *Pseudomonas aeruginosa* and PBP1B^Pa^). We set up reactions in the presence or absence of a soluble version of the lipoprotein activator LpoP^Pa^(sol) for PBP1B^Pa^ (Greene *et al*., 2018). There is currently no reported activator of PBP1B^Ab^, but next to the gene encoding PBP1B^Ab^ we identified a hypothetical gene encoding a lipoprotein containing two tetratricopeptide repeats (Uniprot code D0C5L6) (Figure 2 – figure supplement 4) which we subsequently found to activate PBP1B^Ab^ (see below, Figure 2 – figure supplement 5). We named this protein LpoP^Ab^ and purified a version without its lipid anchor, called LpoP^Ab^(sol). We were able to monitor PG synthesis activity by FRET for both PBPs in the presence or absence of their (hypothetical) activators, using the Triton X-100-solubilized (Figure 2 – figure supplements 6 and 7) or liposome-reconstituted proteins (Figure 2E-H, Figure 2 – figure supplement 2C-D). Our experiments revealed differences in the activities and effect of activators between both PBP1B-homolgoues which we discuss in the following paragraphs.

PBP1B^Ab^ showed GTase activity in the presence of Triton X-100 (Figure 2 – figure supplement 5A) and was stimulated ∼3.3-fold by LpoP^Ab^(sol) (Figure 2 – figure supplement 5B); LpoP^Ab^(sol) also accelerated the consumption of lipid II-Atto550 and glycan chain polymerization (Figure 2 – figure supplement 5C). We measured a low activity of the detergent-solubilised enzyme in the FRET assay (Figure 2 – figure supplement 6A) and poor production of cross-linked PG (Figure 2 – figure supplement 6C), unlike in the case of the other PBPs. However, the liposome-reconstituted PBP1B^Ab^ displayed a higher TPase activity than the detergent-solubilised enzyme (compare gels on Figure 2E, right panel and Figure 2 – figure supplement 6C). In addition, the final FRET signal was substantially higher in liposomes than in detergents (Figure 2E, Figure 2 – figure supplement 6A). Moenomycin completely blocked FRET development, whilst ampicillin had a negligible effect on the final FRET levels in detergents and only a small effect in liposomes (∼1.2-fold reduction), indicating that intra-chain FRET is the major contributor to FRET (Figure 2E; Figure 2 – figure supplement 6A). LpoP^Ab^(sol) stimulated PBP1B^Ab^, with a higher effect in detergents (5.1-fold increase) than liposomes (∼2.5-fold increase) (Figure 2E, F; Figure 2 – figure supplement 6A-B).

PBP1B^Pa^ displayed robust TPase activity in detergents and liposomes (Figure 2G, right panel; Figure 2 – figure supplement 7C) and ampicillin reduced the final FRET signal by ∼1.8-fold in Triton X-100 and by ∼1.5-fold in liposomes, indicating a substantial contribution of inter-chain FRET to the FRET signal (Figure 2G, Figure 2 – figure supplement 7A). The addition of LpoP^Pa^(sol) resulted in an increase in the final FRET by ∼2.2-fold in the membrane and by ∼2.1-fold in detergents (Figure 2G, Figure 2 – figure supplement 7A), accelerated initial slopes by ∼4.2-fold in the membrane and by ∼11.5-fold in detergents (Figure 2H, Figure 2 – figure supplement 7B); lipid II consumption was increased under both conditions (Figure 2G, right panel; Figure 2 – figure supplement 7C). Overall, these results indicate that LpoP^Pa^(sol) stimulates both GTase and TPase activities in agreement with a recent report (Caveney *et al*., 2020).

### PG synthesis on supported lipid bilayers

As we were able to successfully reconstitute active class A PBPs in membranes and monitor their activity in real time, we next aimed to characterise the behaviour of these enzymes in the membrane in more detail by reconstituting them on supported lipid bilayers (SLBs). SLBs are phospholipid bilayers formed on top of a solid support, usually a glass surface and they allow for studying the spatial organization of transmembrane proteins and their diffusion along the membrane by fluorescence microscopy at high spatio-temporal resolution.

We optimized the reconstitution of PBP1B^Ec^ in SLBs formed with EcPL and used the optimized buffer conditions for activity assays on liposomes. To support lateral diffusion and also improve stability of the proteins incorporated into SLBs, we employed glass surfaces coated with polyethylene glycol (PEG) end-functionalized with a short fatty acid (Roder *et al*., 2011) to anchor the EcPL bilayer (Figure 3A). We noticed a decrease in membrane diffusivity and homogeneity at a high surface density of PBP1B^Ec^ (Figure 3 – figure supplement 1). To prevent disturbing the SLB structure by the inserted protein we reduced the density of PBP1B^Ec^ on SLBs from ∼10^−3^ mol protein/mol lipid in liposomes to a range of 10^−6^ to 10^−5^ mol protein/mol lipid. Using a fluorescently-labelled version of PBP1B^Ec^ reconstituted in SLBs, we were able to track the diffusion of single PBP1B molecules in the plane of lipid membrane in the presence or absence of substrate lipid II by TIRF microscopy (Figure 3B, 3D, Movie 1). PBP1B^Ec^ diffused on these supported bilayers with an average D_coef_ of 0.23±0.06 µm^2^/s. Addition of lipid II slowed down PBP1B^Ec^ diffusion (Figure 3C), resulting in a lower average D_coef_ of 0.10±0.06 µm^2^/s. Upon addition of lipid II, we could not detect a prolonged confined motion within particle tracks (Figure 3D), however the average length of displacements was reduced (Figure 3E). Thus we successfully reconstituted diffusing PBP1B^Ec^ in SLBs and we observed that lipid II-binding slowed down the diffusion of the synthase.

**Figure 3.**
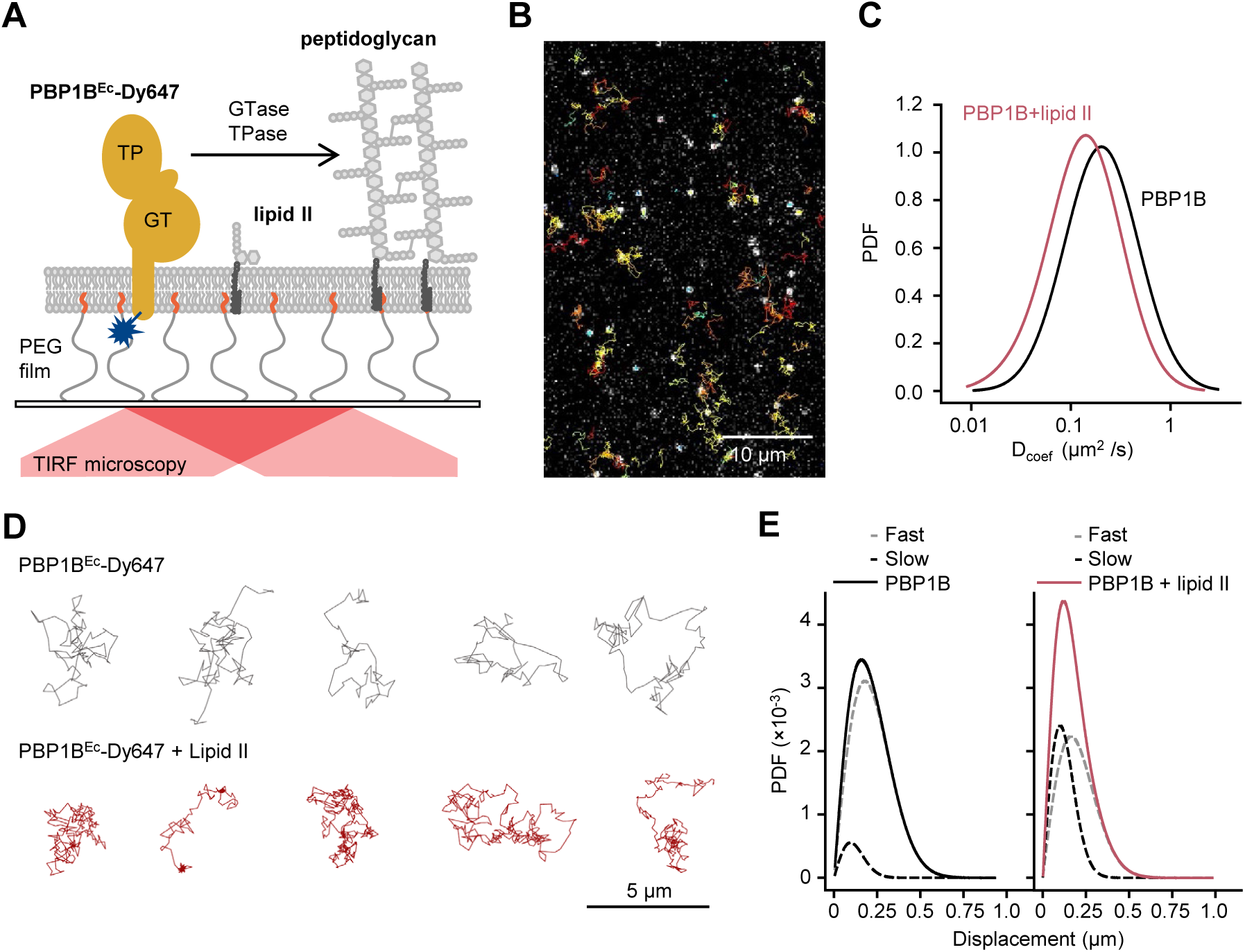
Addition of lipid II slows down diffusion of PBP1B on supported lipid bilayers. (**A**) Schematic illustration of the approach (not to scale). A single-cysteine version of PBP1B^Ec^ (MGC-^64^PBP1B-his C777S C795S) labelled with fluorescent probe Dy647 in its single Cys residue (PBP1B^Ec^-Dy647) was reconstituted into a polymer-supported lipid membrane formed with *E. coli* polar lipids and its diffusion was monitored using TIRF microscopy in the presence or absence of substrate lipid II. (**B**) Single-molecule TIRF micrograph of PBP1B^Ec^-Dy647 diffusing in the lipid membrane in the presence of 1.5 µM lipid II (corresponding to Movie 1). Calculated particle tracks are overlaid. (**C**) Histograms of diffusion coefficients (D_coef_) of PBP1B^Ec^-Dy647 particles in the presence (red) or absence (black) of lipid II. The average D_coef_ decreased from 0.23±0.06 µm^2^/s to 0.1±0.04 µm^2^/s upon addition of lipid II. Values are mean ± SD of tracks from 3 independent experiments. (**D**) Representative tracks for diffusing PBP1B^Ec^-Dy647 particles in the absence (black, top) or presence of lipid II (red, bottom), showing the absence of confined motion in the presence of lipid II. (**E**) Displacement distributions of PBP1B^Ec^-Dy647 particles (solid lines) in the absence (left) or presence (right) of lipid II were analysed using a Rayleigh model incorporating two populations of particles, a fast-diffusing one (grey dashed lines) and a slow-diffusing one (black dashed lines). In the absence of lipid II, only 8±5% of the steps were classified into the slow fraction (121±6nm average displacement), while the majority of steps were of 257±6 nm (fast fraction). The slow fraction increased upon addition of lipid II to 37±5% of the steps, with an average displacement of 132±16 nm.

Next we wanted to confirm that PBP1B^Ec^ remained active to produce planar bilayer-attached PG. We incubated SLBs containing PBP1B^Ec^ with radioactive lipid II and digested any possible PG produced with a muramidase and analysed the digested material by HPLC. Due to the low density and amount of PBP1B^Ec^ on each SLB chamber we expected a small amount of PG product; hence, we included LpoB(sol) to boost the activity of PBP1B^Ec^. Under these conditions about 12% of the added radiolabelled lipid II was incorporated into PG after an overnight incubation (Figure 3 – figure supplement 2A). However, products of both the GTase and TPase activities of PBP1B^Ec^ were detected and these products were absent in the presence of moenomycin (Figure 3 – figure supplement 2B). After overnight PG synthesis reactions with radioactive lipid II, about 32% of the radioactivity remained in the membrane fraction after washing (PG products and unused lipid II) and 68% was in the supernatant. The analysis of the membrane and wash fractions by HPLC (Figure 3 – figure supplement 2C-D) revealed that SLB-reconstituted PBP1B^Ec^ produced crosslinked PG while, importantly, the wash fraction contained no PG products, confirming that the PG synthesis occurred on the SLBs and this PG remained attached to the bilayer. The fraction of membrane-attached radioactivity was almost the same (33%) when PBP1B^Ec^ was not present in the bilayer, indicating that PBP1B^Ec^ did not affect lipid II-binding to the bilayer.

### FRET assay on supported bilayers

Next, we adapted the FRET assay to SLBs and TIRF microscopy taking advantage of the photostability and brightness of the Atto550 and Atto647n probes. Our aim was to visualize PG synthesis by class A PBPs at high resolution as a first step towards understanding PG synthesis at a single molecule level. We used a similar approach as for liposomes, where both Atto550- and Atto647n-labelled lipid II were co-reconstituted with PBP1B^Ec^ on supported lipid bilayers and PG synthesis was triggered by the addition of unlabelled lipid II (Figure 2A). To measure any change in FRET due to PG synthesis, we took advantage of the fact that upon photobleaching of the acceptor probe in a FRET pair, the emitted fluorescence intensity of the donor increases as absorbed energy cannot be quenched by a nearby acceptor (Verveer, 2005, Loose *et al*., 2011). Indeed, we detected an increase in lipid II-Atto550 fluorescence intensity upon photobleaching of the Atto647n probe after the addition unlabelled lipid II and LpoB(sol), indicating the presence of FRET (Figure 4A, Figure 4 – figure supplement 1A). When we bleached the acceptor at different time points of the reaction, we found the FRET signal to increase after a lag phase of ∼8 min. Importantly, there was no FRET increase in the presence of ampicillin (Figure 4B, Figure 4 – figure supplement 1A, Movie 2) or when a GTase-defective PBP1B^Ec^ version (E233Q) was used (Figure 4C). In addition, the FRET signal was abolished when the muramidase cellosyl was added after the PG synthesis reaction (Figure 4C). These results imply that the FRET signal detected by microscopy is primarily due to the transpeptidase activity of PBP1B^Ec^, in agreement with the results obtained on liposomes (Figure 2C).

**Figure 4.**
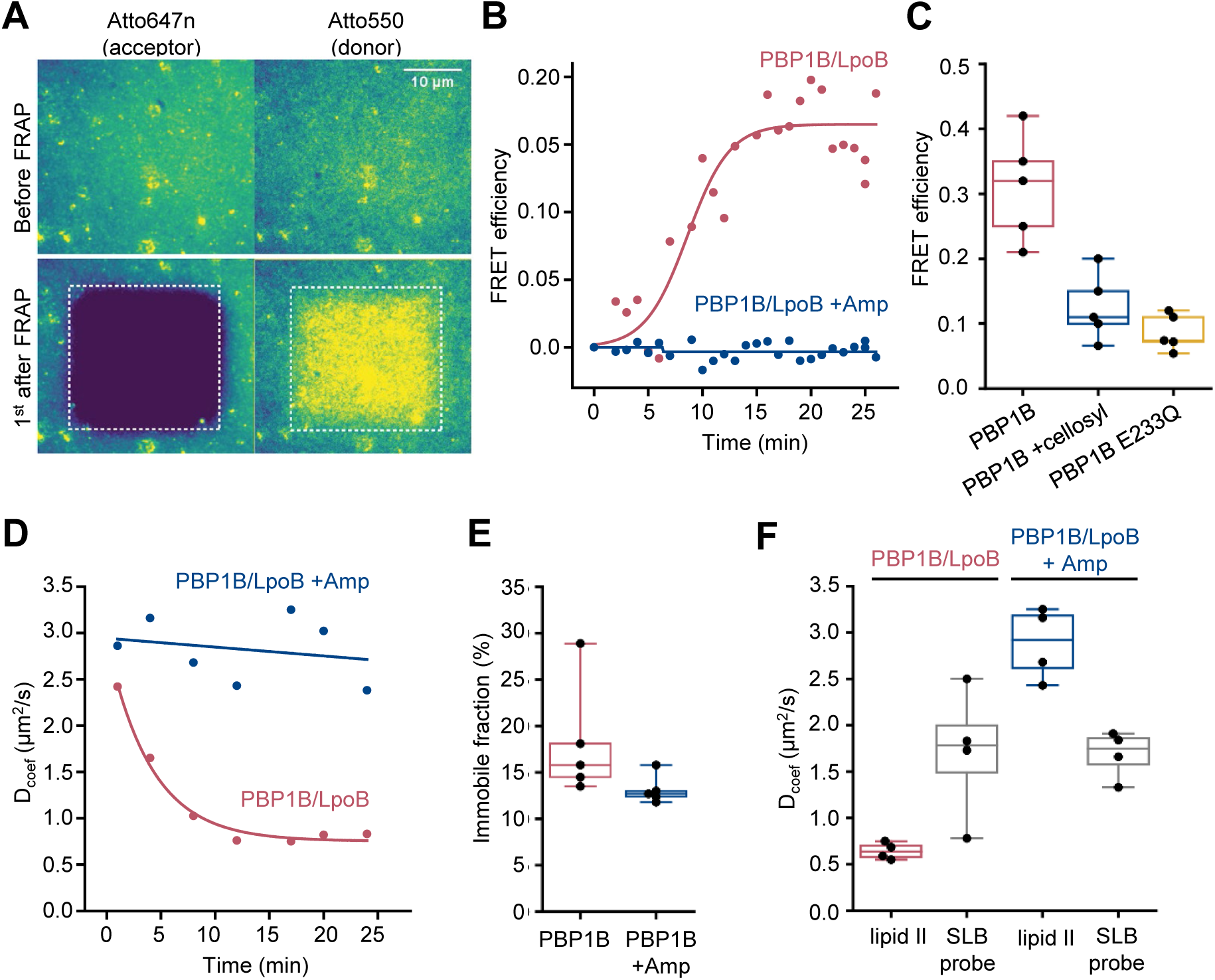
FRET assay on a planar lipid membrane. (**A**) FRET acquisition by TIRF microscopy. PBP1B^Ec^ was reconstituted into a polymer supported lipid membrane to preserve its lateral diffusion. A supported lipid membrane was formed from *E. coli* polar lipid extract supplemented with 0.5 mol% of labelled lipid II (Atto550 and Atto647n at 1:1 ratio). To initiate PG polymerization unlabelled lipid II (10 µM) and of LpoB(sol) (4 µM) were added from the bulk solution. An increase in FRET efficiency was recorded by dual-colour TIRF microscopy: the acceptor (lipid II-Atto647n) was photobleached and the concomitant increase in the donor intensity (lipid II-Atto550) was recorded within a delay of 1 s. (**B**) FRET kinetics of PG polymerization and cross-linking. Inhibition of PBP1B^Ec^ TPase activity with 1 mM ampicillin did not produce any changes in the donor intensity, confirming that FRET signal is specific to cross-linked PG. A sigmoid (straight lines) was fitted to the data to visualise the lag in the increase of FRET signal. (**C**) FRET efficiency was measured after a round of PG synthesis before and after digestion with the muramidase cellosyl. After cellosyl digestion, FRET efficiency decreased by 2.5-fold, resulting in a FRET signal comparable to the one of a control surface with a GTase-defective PBP1B^Ec^(E233Q), performed in parallel. Each dot corresponds to a different surface area within the same sample. (**D**) Quantification of the diffusion coefficient of lipid II-Atto647n over the time course of PG polymerization (left panel) from the experiment presented in **B**, calculated from the dynamics of the recovery of lipid II-Atto647n signal within the photobleached ROI. (**E**) Quantification of the fraction of immobile lipid II-Atto647n from several experiments as the one depicted in **B**, each dot represents the value from a different experiment. (**F**) Diffusion of lipid II-Atto647n or a phospholipid bound probe labelled with Alexa 488 (SLB) was recorded in a FRAP assay, using a 1 s delay and dual-colour imaging, 30 min after initiation of PG synthesis by addition of lipid II and LpoB(sol). Only the diffusion of lipid II, but not of a fluorescently labelled, His_6_-tagged peptide attached to dioctadecylamine-tris-Ni^2+^-NTA, was affected by the presence of ampicillin during the PG synthesis reaction.

### PG synthesised on supported lipid bilayers

As our experiments confirmed that the PG synthesized by PBP1B^Ec^ on SLBs remained attached to the bilayer, we next analysed the lateral diffusion of lipid II-Atto647n and its products during PG synthesis reactions. We first analysed the recovery of fluorescence intensity after photobleaching to monitor the diffusion of lipid II-Atto647n during PG synthesis (Figure 4D). Only when crosslinking was permitted (absence of ampicillin), the diffusion coefficient of lipid II-Atto647n decreased 2 to 3-fold in a time-dependent manner. The time needed to reach the minimum diffusivity value (∼10 min) was similar to the lag detected in the increase of FRET efficiency (Figure 4B). The fraction of immobile lipid II-Atto647n did not change significantly in the presence or absence of ampicillin (13% ± 2% or 18% ± 6%, respectively, p-value = 0.15) (Figure 4E), indicating that the crosslinked PG was still mobile under these conditions, but diffused more slowly. We also compared the diffusion of lipid II-Atto647n during the PG synthesis reaction with that of an AlexaFluor 488-labelled membrane-anchored peptide in the presence or absence of ampicillin (Figure 4F, Figure 4 – figure supplement 2B). The inhibition of TPase by ampicillin only affected the diffusivity of lipid II (2.9 ± 0.4µm^2^/s with ampicillin and 0.67 ± 0.1 µm^2^/s without), while that of the lipid probe remained unchanged (1.6 ± 0.65 µm^2^/s with ampicillin and 1.94 ± 0.62 µm^2^/s without). This shows that the membrane fluidity was not altered by the PG synthesis reaction and therefore was not the cause of the change in lipid II diffusivity upon transpeptidation. As the immobile fraction of labelled lipid II did not increase after PG synthesis and the diffusion was reduced only 2 to 3-fold, we concluded that lipid II-Atto647n was incorporated into small groups of crosslinked glycan chains which can still diffuse on the bilayer.

In summary, we report the incorporation of active PBP1B^Ec^ into supported lipid bilayers, where we could track a decrease in the diffusion of the protein and its substrate during PG synthesis reactions. Using this system we detected an increase in FRET upon initiation of PG synthesis, only occurring when transpeptidation was not inhibited.

## DISCUSSION

Even though class A PBPs are membrane proteins and PG precursor lipid II is embedded in the bilayer, few studies have provided information about the activity of these important enzymes in a membrane environment. Here we developed a new assay that reports on PG synthesis by these enzymes in detergents, on liposomes or on supported lipid bilayers.

### Intra-chain *vs* inter-chain FRET

For all PBPs and conditions tested, FRET increased when only the GTase domain was active (i.e. when FRET occurred between probes incorporated along the same strand), but the FRET signal was always higher when transpeptidase was active (Figures 1, 2 and Figure 2 – figure supplements 6 and 7). For detergent-solubilised PBP1B^Ec^, the FRET curve closely followed the rate of the production of cross-linked PG as determined by HPLC analysis of the products (Figure 1E-G). These results suggest that inter-chain FRET (arising from both fluorophores present on different, adjacent glycan chains) was a main component of the total FRET signal. Why is this the case? FRET depends on the distance and orientation of the two probes. It might be sterically unfavourable that two large Atto550 and Atto647n containing lipid II molecules simultaneously occupy the donor and acceptor sites in the GTase domain (van&t Veer, 2016), preventing the incorporation of probes (and high FRET) at successive subunits on a single glycan chain. Indeed, for all PBPs tested either in detergents or liposomes, the incorporation of labelled lipid II into glycan chains was more efficient when unlabelled lipid II was present and for most enzymes an activator was required to polymerize glycan chains using labelled lipid II in the absence of unlabelled lipid II. We thus hypothesize that the TPase activity brings glycan chains to close proximity, reducing the distance between probes sufficiently to produce high levels of FRET (Figure 5).

**Figure 5.**
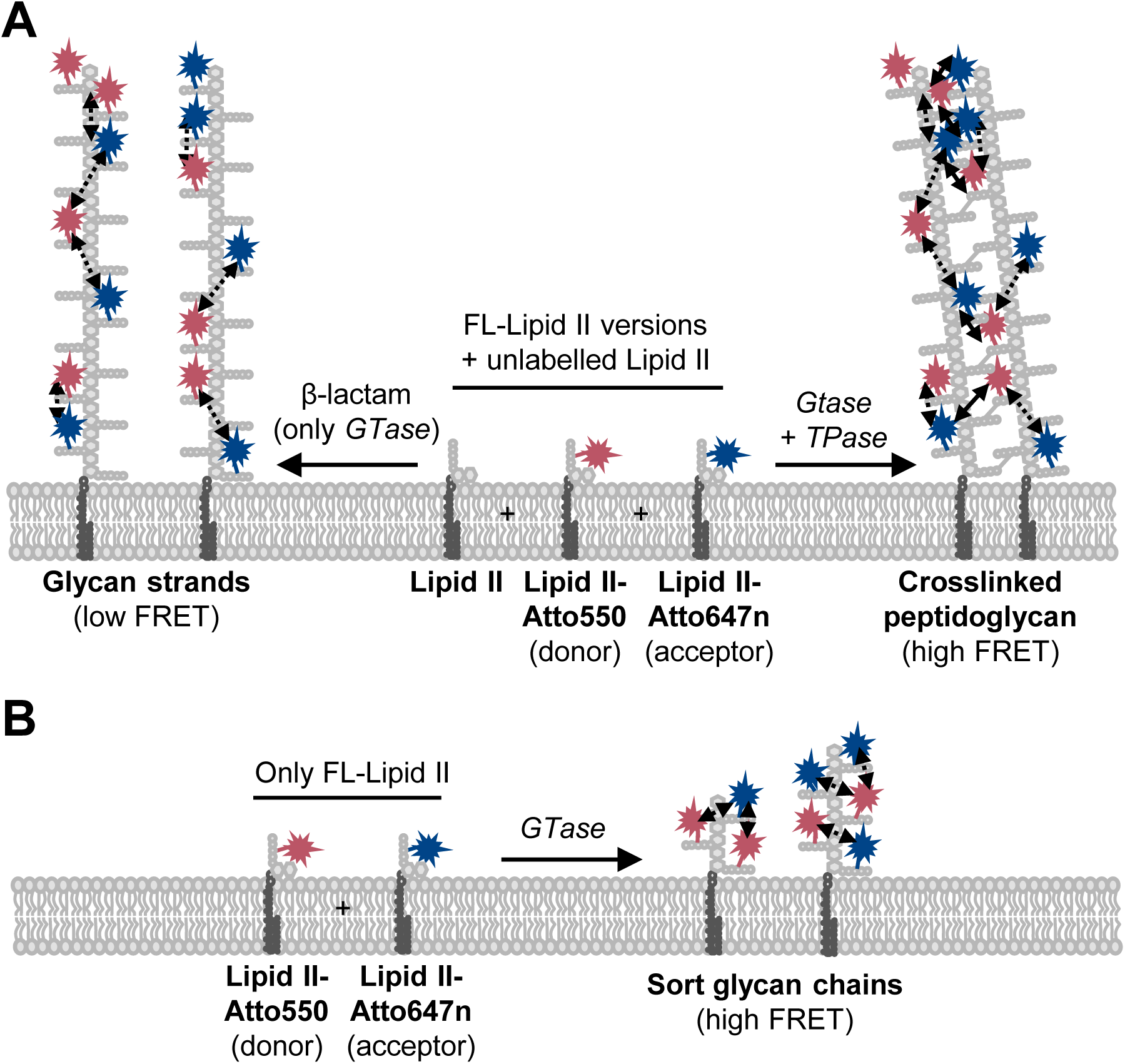
PG synthesis with labelled lipid II versions and detection of FRET. (**A**) A mixture of Atto550-lipid II, Atto647n-lipid II and unlabelled lipid II is utilized by a class A PBP with or without inhibition of the TPase activity by a β-lactam. FRET can only occur between fluorophores within the same glycan strand in linear glycan chains produced in the presence of a β-lactam (left reaction, dashed arrows). When the TPase is active (right reaction) FRET can occur either between probes within the same strand (dashed arrows) or between probes on different strands of the cross-linked PG product (solid arrows). We hypothesize that at any time only one labelled lipid II molecule occupies the two binding sites in the GTase domain and that therefore two probes within the same strand are separated by at least one subunit. As a result, average distances between probes in different strands may be shorter than between probes within the same strand and thus inter-chain FRET contributes stronger to the total FRET signal than intra-chain FRET. (**B**) Lipoprotein-stimulated PBPs produced short chains when labelled lipid II versions were incubated in the absence of unlabelled lipid II (e.g., Figure 1B and Figure 1-figure supplement 1C). In this situation crosslinking does not occur due to the attachment of the probe to the mDAP residue in the pentapeptide. Within these short strands intra-chain FRET is stronger than within the long glycan strands depicted in (**A**), due to a shorter average distance between the probes.

### Coupled reactions in class A PBs

An elegant recent report (Catherwood *et al*., 2020) described the use of a coupled D-Ala release assay to determine the kinetic parameters of the TPase activity of PBP1B^Ec^ against different substrates and this study also confirmed the previously reported activation of the TPase of PBP1B by LpoB in the presence of detergents (Egan *et al*., 2014, Egan *et al*., 2018, Lupoli *et al*., 2014). The authors of the recent report (Catherwood *et al*., 2020) discussed that the LpoB-mediated TPase activation explains the essentiality of LpoB for PBP1B function in the cell. However, this view ignores previously published data demonstrating that the essentiality of LpoB can be readily explained by its primary effect, the >10-fold stimulation of PBP1B’s GTase (Egan *et al*., 2014). TPase reactions follow and depend on ongoing GTase reactions (Bertsche *et al*., 2005). Interestingly, LpoB-activated PBP1B produces a hyper-crosslinked PG (Typas *et al*., 2010, Egan *et al*., 2018), suggesting that LpoB stimulates the TPase more than the GTase. In the cell, a protein associated with the Tol system, CpoB, modulates this hyper-stimulation of the TPase when coordinating outer membrane constriction and PG synthesis during cell division (Gray *et al*., 2015). The observed stimulation of both reactions by LpoB is consistent with conformational changes in the regulatory UB2H domain in PBP1B that occur upon LpoB binding and that affect amino acid residues pointing towards both domains (Egan *et al*., 2018). A limitation of the recent kinetic study is that authors used assay conditions (e.g. very low enzyme concentration) at which PBP1B^Ec^ is virtually inactive without an activator (Catherwood *et al*., 2020) as demonstrated previously (Pazos *et al*., 2018, Muller *et al*., 2007), thus the study likely substantially overestimated the extent of TPase activation by LpoB.

*P. aeruginosa* uses a structurally different lipoprotein activator, LpoP, to stimulate its PBP1B (Greene *et al*., 2018). Here, we identified an LpoP homologue in *A. baumannii* and showed that both, LpoP^Ab^ and LpoP^Pa^ significantly activated their cognate PBP1B. Interestingly, LpoP^Ab^ stimulated the GTase and not TPase of PBP1B^Ab^ while LpoP^Pa^ stimulated both activities in PBP1B^Pa^ which may illustrate how different species have tailored their activators to their specific needs. Importantly, PBP1B^Ab^ TPase activity was higher in liposomes than in detergents, which serves as a reminder that detergents are not always neutral solubilising agents and they can affect the activity of membrane proteins.

### Towards single-molecule PG synthesis

We also adapted the FRET assay to supported lipid bilayers and super resolution microscopy to study how PBP1B^Ec^ polymerizes PG on SLBs (Figure 4). As with the liposome assays, we detected an increase in FRET signal upon triggering PG synthesis that correlated with transpeptidation. Importantly we could follow the diffusion of the reaction products, which indicates that PBP1B^Ec^ does not completely cover the surfaces with a layer of PG but instead produced smaller patches of cross-linked glycan chains. We attribute this to the fact that PBP1B^Ec^ was reconstituted at a very low density in order to ensure the homogeneity and stability of the SLBs. Remarkably, we detected a reduction of PBP1B^Ec^ diffusivity in the presence of lipid II (Figure 3). Previous *in vivo* single-molecule tracking of fluorescent-protein tagged class A PBPs reported the presence of two populations of molecules, a fast diffusing one and an almost immobile one with a near-zero diffusing rate which was assumed to be the active population (Cho *et al*., 2016, Lee *et al*., 2016, Vigouroux *et al*., 2020). Our result supports this interpretation, although more experiments are required to further explore this point.

Several real time methods to study PG synthesis *in vitro* are described in the literature. However, most of these report on either the GTase or TPase reaction, but not both at the same time, and most available methods are not applicable to the membrane. The scintillation proximity assay by Kumar et al. reports on PG production in a membrane environment and in real time, but it is rather crude in that it uses membrane extract instead of purified protein and relies on the presence of lipid II synthesizing enzymes present in the extract (Kumar *et al*., 2014). Moreover, it is uses radioactivity detection and is not amenable to microscopy, in contrast to methods based on fluorescently-labelled substrates. An important advantage of our new assay over other real-time TPase assays is that it uses natural substrates for transpeptidation,, i.e. nascent glycan strands, instead of mimics of the pentapeptide, and its ability to measure the activities in a natural lipid environment.

Our new FRET assay can potentially be adopted to assay PG synthases in the presence of interacting proteins, for example monofunctional class B PBPs in the presence of monofunctional GTases (cognate SEDS proteins or Mtg proteins) or interacting class A PBPs (Meeske *et al*., 2016, Bertsche *et al*., 2006, Sjodt *et al*., 2020, Derouaux *et al*., 2008, Banzhaf *et al*., 2012, Sjodt *et al*., 2018). In addition, our assay has the potential to be adopted to high throughput screening for new antimicrobials.

## MATERIALS AND METHODS

### Chemicals

[^14^C]Glc*N*Ac-labelled lipid II and the lysine or *m*DAP forms of lipid II were prepared as published (Breukink *et al*., 2003, Bertsche *et al*., 2005). Lipid II-Atto550 and Lipid II-Atto647n were prepared from the lysine form of lipid II, and Atto550-alkyne or Atto647n-alkyne (Atto tec, Germany) as previously described (Mohammadi *et al*., 2014, Egan *et al*., 2015). Polar lipid extract from *E. coli* (EcPL), 1,2-dipalmitoleoyl-*sn*-glycero-3-phosphocholine (DOPC), 1-palmitoyl-2-oleoyl-*sn*-glycero-3-phospho-(1&-*rac*-glycerol) (POPG) and tetraoleoyl cardiolipin (TOCL) were obtained from Avanti Polar Lipids (USA). Lipids were resuspended in chloroform:methanol (2:1) at a concentration of 20 g/L, aliquoted and stored at -20°C. Triton X-100, ampicillin, phenylmethylsulfonyl fluoride (PMSF), protease inhibitor cocktail (PIC) and β-mercaptoethanol were from Merck. n-dodecyl-beta-D-maltopyranoside (DDM) was purchased from Anatrace (USA). Moenomycin was purchased from Hoechst, Germany. All other chemicals were from Merck.

### Cloning

#### Construction of overexpression vector pKPWV1B

The plasmid pKPWV1B was constructed for overexpression of full-length *A. baumannii* PBP1B (PBP1B^*Ab*^: aa 1-798) with a cleavable N-terminal oligo-histidine tag (His_6_ tag). Therefore, the gene *mrcB* was amplified using the Phusion high fidelity DNA polymerase and the oligonucleotides PBP1B.Acineto-NdeI_f and PBP1B.Acineto-BamHI_r and genomic DNA of *A. baumannii* 19606 (ATCC) as template. The resulting PCR fragment and the Plasmid DNA of the overexpression vector pET28a(+) (Novagen) were digested with *Nde*I and *BamH*I, ligated and transformed into chemical competent *E. coli* DH5α cells with kanamycin selection. Plasmid DNA of transformants was isolated and send for sequencing using following oligonucleotides: Seq1_rev_PBP1B_Acineto, Seq2_fwd_PBP1B_Acineto, Seq3_fwd_PBP1B_Acineto, Seq4_fwd_PBP1B_Acineto.

#### Construction of overexpression vector pKPWVLpoP

The sequence of the hypothetical PBP1B activator of *Acinetobacter baumannii* 19606 (LpoP^*Ab*^: NCBI reference number: WP_000913437.1) contains a TPR fold and was found by blast analysis through its homology to *Pseudomonas aeruginosa* LpoP (30% identity). The plasmid pKPWVLpoP was purchased from the company GenScript. The gene was synthesized without the first 51 nucleotides (encoding the 17 amino acids of the signal peptide) and with codon optimization for overexpression in *Escherichia coli*. The codon optimized gene was subcloned in the overexpression vector pET28a(+) using the cloning sites *Nde*I and *BamH*I enabling the overexpression of the protein with an N-terminal oligo-histidine tag.

#### MGC-^64^PBP1B-his C777S/C795S

This fusion protein contains PBP1B with the substitution of the N-terminal cytoplasmic tail for residues MGC and the addition of a hexahistine tag at the C-terminus. To obtain this construct, the regions coding for aminoacids 64 to 844 of PBP1B were amplified from genomic DNA using oligonucleotides PBP1B-MGC-F and PBP1B-CtermH-R. The resulting product was cloned into pET28a+ vector (EMD Biosciences) after digestion with NcoI and XhoI. C777S and C795S mutations were introduced using the QuikChange Lightning mutagenesis kit (Agilent) using oligonucleotide primers C777S-D, C777S-C, C795S-D and C795S-C The resulting plasmid was called pMGCPBP1BCS1CS2.

### Purification and labelling of proteins

The following proteins were purified following published protocols: PBP1B^Ec^ (Bertsche *et al*., 2006), LpoB(sol) (Egan *et al*., 2014), PBP1B^Pa^ (Caveney *et al*., 2020), LpoP^Pa^(sol) (Caveney *et al*., 2020). All chromatographic steps were performed using an AKTA PrimePlus system (GE Healthcare).

#### E. coli PBP1B

The protein was expressed as a fusion with an N-terminal hexahistidine tag in *E. coli* BL21(DE3) pDML924 grown in 4 L of autoinduction medium (LB medium supplemented with 0.5% glycerol, 0.05% glucose, and 0.2% α-lactose) containing kanamycin at 30 °C for ∼16h. Cells were harvested by centrifugation (10,000 × g, 15 min, 4 °C) and the pellet resuspended in 80 mL of buffer I (25 mM Tris-HCl, 1 M NaCl, 1 mM EGTA, 10% glycerol, pH 7.5) supplemented with 1× protease inhibitor cocktail (PIC, Sigma-Aldrich), 100 µM phenylmethylsulfonyl fluoride (PMSF, Sigma-Aldrich) and DNase I. After disruption by sonication on ice, membrane fraction was pelleted by centrifugation (130,000 × g for 1 h at 4 °C) and resuspended in buffer II (25 mM Tris-HCl, 1 M NaCl, 10% glycerol, 2% Triton X-100, pH 7.5) by stirring at 4 °C for 24 h. Extracted membranes were separated from insoluble debris by centrifugation (130,000 × g for 1 h at 4 °C) and incubated for 2h with 4 mL of Ni^2+^-NTA beads (Novagen) equilibrated in buffer III (25 mM Tris-HCl, 1 M NaCl, 20 mM imidazole, 10% glycerol, pH 7.5). Beads were washed 10 times with 10 mL of buffer III and the protein was eluted with 3 mL buffer IV (25 mM Tris-HCl, 0.5 M NaCl, 20 mM imidazole, 10% glycerol, pH 7.5). His-PBP1B containing fractions were pooled and treated with 2 U/mL of thrombin (Novagen) for 20 h at 4 °C during dialysis against dialysis buffer I (25 mM Tris-HCl, 0.5 M NaCl, 10% glycerol, pH 7.5). Protein was then dialysed in preparation for ion exchange chromatography, first against dialysis buffer II (20 mM sodium acetate, 0.5 M NaCl, 10% glycerol, pH 5.0); then against dialysis buffer II with 300 mM NaCl; and finally against dialysis buffer II with 100 mM NaCl. Finally, the sample was applied to a 1 mL HiTrap SP column (GE Healthcare) equilibrated in buffer A (20 mM sodium acetate, 100 mM NaCl, 10% glycerol, 0.05% reduced Triton X-100, pH 5.0). The protein was eluted with a gradient from 0 to 100% buffer B (as A, with 2 M NaCl) over 14 mL PBP1B-containing fractions were pooled and dialysed against storage buffer (20 mM sodium acetate, 500 mM NaCl, 10% glycerol, pH 5.0) and stored at –80 °C.

#### A. baumannii 19606 PBP1B

The protein was expressed in *E. coli* BL21 (DE3) freshly transformed with plasmid pKPWV1B using the same protocol as PBP1B^Ec^. Cells were harvested by centrifugation (6,200 × *g* for 15 min at 4 °C) and resuspended in 120 mL of PBP1B^*Ab*^ buffer I (20 mM NaOH/H_3_PO_4_, 1 M NaCl, 1 mM EGTA, pH 6.0) supplemented with DNase I, PIC (1:1,000 dilution) and 100 µM PMSF. After disruption by sonication on ice, the membrane fraction was pelleted by centrifugation (130,000 × *g* for 1 h at 4 °C) and resuspended in PBP1B^*Ab*^ extraction buffer (20 mM NaOH/H_3_PO_4_, 1 M NaCl, 10% glycerol, 2% Triton X-100, pH 6.0) supplemented with PIC and PMSF by stirring at 4 °C for 16 h. Extracted membranes were separated from insoluble debris by centrifugation (130,000 × g for 1 h at 4 °C) and incubated with 4 mL of Ni^2+^-NTA beads equilibrated in PBP1B^*Ab*^ extraction buffer containing 15 mM imidazole. Beads were washed 10 times with 10 mL of PBP1B^*Ab*^ wash buffer (20 mM NaOH/H_3_PO_4_, 10% Glycerol, 0.2% Triton X-100, 1M NaCl, 15 mM Imidazole, pH 6.0) and the protein was eluted with 3 mL buffer IV PBP1B^*Ab*^ elution buffer (20 mM NaOH/H_3_PO_4_, 10% Glycerol, 0.2% Triton X-100, 1 M NaCl, 400 mM Imidazole, pH 6.0).

PBP1B^*Ab*^-containing fractions were pooled and dialyzed in preparation for ion exchange chromatography, first against PBP1B^*Ab*^ dialysis buffer I (20 mM sodium acetate, 1 M NaCl, 10% glycerol, pH 5.0), then against PBP1B^*Ab*^ dialysis buffer II (20 mM sodium acetate, 300 mM NaCl, 10% glycerol, pH 5.0) and finally against PBP1B^*Ab*^ dialysis buffer III (10 mM sodium acetate, 100 mM NaCl, 10% glycerol, pH 5.0). The sample was centrifuged for 1 h at 130,000 × *g* and 4 °C and the supernatant was applied to a 5 mL HiTrap SP HP column equilibrated in PBP1B^*Ab*^ buffer A (20 mM sodium acetate, 100 mM NaCl, 10% glycerol, 0.2% Triton X-100, pH 5.0). The protein was eluted from 0 to 100% PBP1B^*Ab*^ buffer B (20 mM sodium acetate, 2 M NaCl, 10% glycerol, 0.2% Triton X-100, pH 5.0) over 70 mL. PBP1B^Ab^-containing fractions were pooled and dialysed against PBP1B^*Ab*^ storage buffer (10 mM sodium acetate, 500 mM NaCl, 0.2% Triton X-100, 20% glycerol, pH 5.0) and stored at -80 °C.

#### P. aeruginosa PBP1B

The protein was expressed on *E. coli* BL21(DE3) freshly transformed with plasmid pAJFE52 which encodes PBP1BPa as a fusion with an N-terminal hexahistidine tag in *E. coli* BL21(DE3). Cells were grown in 4 L of LB at 30 °C and expression was induced for 3 h with 1 mM IPTG when the culture reached an OD_578_ of 0.6. PBP1B^Pa^ was extracted and purified using the same protocol as for *E. coli* PBP1B with the exception that only 2 mL of Ni^2+^ beads were used.

#### MGC-^64^PBP1B-his C777S/C795S

This protein was expressed in *E. coli* BL21(DE3) freshly transformed with plasmid pMGCPBP1BCS1CS2 and subsequently purified using the same protocol as for the WT protein, except for the addition of 1 mM TCEP to all purification buffers. The protein was labelled with Dy647-maleimide probe (Dyomics, Germany) following instructions from the manufacturer. Briefly, 10.2 µM protein was incubated with 100 µM probe and 0.5 mM TCEP for ∼20 h at 4 °C and free probe was removed by desalting using a 5 mL HiTrap desalting column (GE Healthcare).

#### LpoB(sol)

The protein was expressed on *E. coli* BL21(DE3) transformed with pET28His-LpoB(sol). Cells were grown in 1.5 L of LB plus kanamycin at 30 °C to an OD_578_ of 0.4–0.6 and expression was induced with 1 mM of IPTG for 3 h at 30 °C. Cells were pelleted and resuspended in buffer I (25 mM Tris-HCl, 10 mM MgCl_2_, 500 mM NaCl, 20 mM imidazole, 10% glycerol, pH 7.5) plus DNase, PIC and PMSF. Cells were disrupted by sonication on ice and centrifuged (130,000 × g, 1 h, 4 °C) to remove debris. The supernatant was applied to a 5 mL HisTrap HP column (GE Healthcare) equilibrated in buffer I. After washing with buffer I, the protein was eluted with a stepwise gradient with buffer II (25 mM Tris-HCl, 10 mM MgCl_2_, 500 mM NaCl, 400 mM imidazole, 10% glycerol, pH 7.5). Fractions containing the protein were pooled and the His-tag was removed by addition of 2 U/mL of thrombin while dialysing against buffer IEX-A (20 mM Tris-HCl, 1000 mM NaCl, 10% glycerol, pH 8.3). Digested protein was applied to a 5 mL HiTrap Q HP column (GE Healthcare) at 0.5 mL/min. LpoB(sol) was collected in the flow through, concentrated and applied to size exclusion on a Superdex200 HiLoad 16/600 column (GE Healthcare) at 1 mL/min in a buffer containing 25 mM HEPES-NaOH, 1 M NaCl, 10% glycerol at pH 7.5. Finally, the protein was dialysed against storage buffer (25 mM HEPES-NaOH, 200 mM NaCl, 10% glycerol at pH 7.5) and stored at –80 °C.

#### A. baumannii 19606 LpoP(sol)

The protein was expressed on *E. coli* BL21(DE3) transformed with plasmid pKPWVLpoP. Cells were grown over night at 30 °C in 4 L of autoinduction medium. Cells were pelleted by centrifugation (6,200 × *g* for 15 min at 4 °C) and resuspended in 80 mL of buffer I (25 mM Tris/HCl, 10 mM MgCl_2_, 1 M NaCl, 20 mM Imidazole, pH 7.5) supplemented with DNase I, PIC (1:1,000 dilution) and 100 µM PMSF. Cells were disrupted by sonication on ice and centrifuged (130,000 × *g* for 1 h at and 4 °C) to removed debris. The supernatant was incubated for 1h with 6 mL Ni-NTA beads preequilibrated in buffer I at 4 °C with gentle stirring. The resin was split in 2 columns, each washed 10 times with 5 mL wash buffer (25 mM Tris/HCl, 10 mM MgCl_2_, 1 M NaCl, 20 mM Imidazole, pH 7.5) and the protein was eluted 7 times with 2 mL of elution buffer (25 mM Tris/HCl, 10 mM MgCl_2_, 1 M NaCl, 400 mM Imidazole, pH 7.5). The best fractions according to SDS-PAGE analysis were pooled and dialyzed stepwise against increasing percentage of dialysis buffer I (25 mM HEPES/NaOH, 10 mM MgCl_2_, 200 mM NaCl, 10% glycerol, pH 7.5). Thrombin (9 units) was added to the protein to cleave the N-terminal His_6_ tag over night at 4 °C. The successful cleavage of the N-terminal His_6_ tag was confirmed by SDS-PAGE. The protein was diluted 2× with 25 mM HEPES/NaOH, 10 mM MgCl_2_, 10% glycerol, pH 7.5 to reduce the amount of NaCl down to 100 mM. The protein was applied to a 5 mL HiTrap SP HP column and washed with buffer A (25 mM HEPES/NaOH, 10 mM MgCl_2_, 100 mM NaCl, 10% glycerol, pH 7.5). The protein was then eluted with a gradient of 100 mM to 1 M NaCl over 50 mL at 1 mL/min using increasing percentage of buffer B (25 mM HEPES/NaOH, 10 mM MgCl_2_, 1 M NaCl, 10% glycerol, pH 7.5). Fractions were collected and analysed by SDS-PAGE. The best fractions were pooled, dialysed against 25 mM HEPES/NaOH, 200 mM NaCl, 10% Glycerol, 10 mM MgCl_2_, pH 7.5 and the protein were stored at -80 °C.

#### P. aeruginosa LpoP(sol)

The protein was expressed on *E. coli* BL21(DE3) freshly transformed with from plasmid pAJFE57, encoding His_6_-LpoP^Pa^(sol). Cells were grown on 1.5 L LB at 30°C to an OD_578_ of 0.5 and expression was induced for 3h by addition of 1 mM IPTG. After harvesting, cells were resuspended in 80 mL of 25 mM Tris-HCl, 500 mM NaCl, 20 mM imidazole, 10% glycerol at pH 7.5. After addition of PIC and 100 µM PMSF, cells were disrupted by sonication on ice. Debris was removed by centrifugation (130,000 × g, 1 h, 4 °C) and the supernatant was applied to a 5 mL HisTrap column equilibrated in resuspension buffer. After washing with 25 mM Tris-HCl, 1 M NaCl, 40 mM imidazole, 10% glycerol at pH 7.5, protein was eluted with 25 mM Tris-HCl, 500 mM NaCl, 400 mM imidazole, 10% glycerol at pH 7.5. Fractions containing His-LpoP^Pa^(sol) were pooled and the His-tag was removed by addition of 4 U/mL of thrombin while dialysing against 20 mM Tris-HCl, 200 mM NaCl, 10% glycerol at pH 7.5 for 20 h at 4 °C. The sample was concentrated and further purified by size exclusion column chromatography at 0.8 mL/min using a HiLoad 16/600 Superdex 200 column equilibrated in 20 mM Hepes-NaOH, 200 mM NaCl, 10% glycerol at pH 7.5. LpoP^Pa^-containing fractions were pooled, concentrated, aliquoted and stored at -80°C.

### PG synthesis assays in the presence of detergents

#### In vitro peptidoglycan synthesis assay using radiolabelled lipid II in detergents

To assay the *in vitro* PG synthesis activity of PBP1B^Ec^ with radiolabelled lipid II substrate in the presence of detergent we used a previously published assay (Banzhaf *et al*., 2012, Biboy *et al*., 2013). Final reactions included 10 mM HEPES/NaOH pH 7.5, 150 mM NaCl, 10 mM MgCl_2_ and 0.05 % Triton X-100. The concentration of PBP1B^Ec^ was 0.5 µM. Reactions were carried out for 1 h at 37°C. Reactions were stopped by boiling for 5 min. Digestion with cellosyl, reduction with sodium borohydride and analysis by HPLC were performed as described (Biboy *et al*., 2013).

#### FRET-based in vitro peptidoglycan synthesis assay in detergents

For assays in detergents, samples contained 50 mM HEPES/NaOH pH 7.5, 150 mM NaCl, 10 mM MgCl_2_, and 0.05% Triton X-100 in a final volume of 50 µL. PBP1B^Ec^, PBP1B^Ab^ or PBP1B^Pa^ were added at a concentration of 0.5 µM. When indicated, activators LpoB(sol), or LpoP^Ab^(sol), or LpoP^Pa^(sol) were added at a concentration of 2 µM. Reactions were started by the addition of an equimolar mix of lipid II, lipid II-Atto550 and lipid II-Atto647n, each at 5 µM and monitored by measuring fluorescence using a Clariostar plate reader (BMG Labtech, Germany) with excitation at 540 nm and emission measurements at 590 nm and 680 nm. Reactions were incubated at the indicated temperature for 60 or 90 min. After the reaction emission spectra from 550 to 740 nm were taken in the same plate reader with excitation at 522 nm. When indicated ampicillin was added at 1 mM and moenomycin was added at 50 µM. After plate reader measurements, reactions were stopped by boiling for 5 min, vacuum-dried using a speed-vac desiccator and analysed by Tris-Tricine SDS-PAGE as previously described (Van&t Veer *et al*., 2016).

FRET reactions in the presence of radiolabelled lipid II described in Figure 1E-F were performed using the same buffer and substrate and enzyme concentrations as for the plate reader assay but in a final volume of 350 µL. Samples were incubated at 25 °C with shaking using an Eppendorf Thermomixer. 50 µL aliquots were taken out at the indicated times and reactions were stopped by addition of 100 µM moenomycin. Samples were then transferred to a 96-well plate to measure FRET as described above. Finally, samples were transferred back to Eppendorf tubes, digested with cellosyl and reduced with sodium borohydride as previously described (Biboy *et al*., 2013).

#### Continuous glycosyltransferase (GTase) assay using dansylated lipid II

Continuous fluorescence GTase assays using dansylated lipid II and *A. baumannii* PBP1B were performed as previously described (Schwartz *et al*., 2001, Offant *et al*., 2010, Egan & Vollmer, 2016). Samples contained 50 mM HEPES/NaOH pH 7.5, 105 mM NaCl, 25 mM MgCl_2_, 0.039% Triton X-100 and 0.14 µg/µL cellosyl muramidase in a final volume of 60 µL. PBP1B^Ab^ was added at a concentration of 0.5 µM. When indicated, LpoP^Ab^(sol) was added at a concentration of 0.5 µM. Reactions were started by addition of dansylated lipid II to a final concentration of 10 µM and monitored by following the decrease in fluorescence over 60 min at 37°C using a FLUOstar OPTIMA plate reader (BMG Labtech, Germany) with excitation at 330 nm and emission at 520 nm. The fold-increase in GTAse was calculated against the mean rate obtained with PBP1B^Ab^ alone at these reaction conditions, at the fastest rate.

#### Time-course GTase assay by SDS-PAGE followed by fluorescence detection

PBP1B^Ab^ at a concentration of 0.5 µM was incubated with 5 µM lipid II-Atto550 and 25 µM unlabelled lipid II in the presence or absence of 1.5 µM LpoP^Ab^(sol). Reactions contained 20 mM HEPES, 150 mM NaCl, 10 mM MgCl_2_, 0.06% TX-100 and 1 mM Ampicillin to block transpeptidation. Aliquots were taken after 0, 2, 5, 10, 30 and 60 min incubation at 37°C, boiled for 10 min to stop reactions and analysed by Tris-Tricine SDS-PAGE followed by fluorescence detection as previously described (Van&t Veer *et al*., 2016).

### PG synthesis in liposomes

#### Reconstitution of class A PBPs in liposomes

Proteoliposomes containing class A PBPs were prepared as described previously with some modifications (Egan *et al*., 2015, Rigaud & Lévy, 2003, Hernández-Rocamora *et al*., 2018). The appropriate lipid or mixture of lipids were dried in a glass test tube under stream of N_2_ to form a lipid film followed by desiccation under vacuum from 2 h. When labelled lipid II was co-reconstituted with the indicated class A PBP, they were added at 1:200 mol:mol phospholipid to each lipid II-Atto550 and lipid II-Atto647n. Resuspension into multilamellar vesicles (MLVs) was achieved by addition of 20 mM Tris/HCl, pH 7.5 with or without 150 mM NaCl as indicated in each experiment and several cycles of vigorous mixing and short incubations in hot tap water. The final lipid concentration was 5 g/L. To form large unilamellar vesicles (LUVs), MLVs were subjected to 10 freeze-thaw cycles and then extruded 10 times through a 0.2 µm filter. LUVs were destabilised by the addition of Triton X-100 to an effective detergent:lipid ratio of 1.40 and mixed with proteins in different protein to lipid molar rations (1:3000 for PBP1B^Ec^ and PBP1B^Pa^, and 1:2000 for PBP1B^Ab^). After incubation at 4°C for 1 h, prewashed adsorbent beads (Biobeads SM2, BioRad, USA; 100 mg per 3 µmol of Triton X-100) were added to the sample to remove detergents. Biobeads were exchanged after 2 and 16 h, followed by incubation with fresh Biobeads for a further 2 h. After removal of Biobeads by short centrifugation at 4,000×*g*, liposomes were pelleted at 250,000×*g* for 30 min at 4°C. The pellet containing proteoliposomes was resuspended using the appropriate buffer. The resuspension was done in a 43% smaller volume than the volume added of lipid II, so that the final concentration of lipids was 11.6 g/L. Samples were then centrifuged for 5 min at 17,000×*g* and 4°C to remove any possible aggregates. The supernatant was then used in the appropriate assays. Liposomes were analysed by SDS-PAGE and, only for liposomes without labelled lipid II, also by bicinchoninic acid assay (Pierce BCA Assay Kit, ThemoFisher Scientific, USA) to determine protein concentration. The concentration of protein for liposomes with labelled lipid II was calculated by densitometry of the samples in SDS-PAGE gels, after reactions were carried out.

#### PBP1B^Ec^ orientation assay

To assess the orientation of liposome-reconstituted PBP1B^Ec^, MGC-^64^PBP1B-his C777S C795S mutant containing a single cysteine in the N-terminal region was reconstituted in liposomes with EcPL as described above. The accessibility of the cysteine was determined using sulfhydryl-reactive fluorescent probe AlexaFluor555-maleimide. Reactions containing 0.5 µM protein, 10 µM AlexaFluor555-maleimide, 0.2 mM TCEP were incubated for 16 h at 4°C in the presence or absence 0.5% Triton X-100.

Reactions were stopped by addition of 5 mM DTT and boiling for 5 min. Samples were loaded in a 10% acrylamide gel and, after electrophoresis, gels were first scanned using an Amersham Typhoon Trio with excitation at 533 nm and a 40 nm-wide band-pass emission filter at 580 nm. The gel was then stained by Coomassie.

#### In vitro peptidoglycan synthesis assay using radiolabelled lipid II in liposomes

The same methodology as in detergents was used to assay the *in vitro* PG synthesis activity of PBP1B^Ec^ in liposomes, with minor modifications. To start reactions, 1.5 nmol [^14^C]-labelled lipid II were dried in a 0.5 mL glass tube using a vacuum concentrator, resuspended in 5 µL of the appropriate liposome buffer, and mixed with liposomes, buffer and MgCl_2_ to a total volume of 50 µL. Final reactions contained 0.5 µM PBP1B^Ec^, 30 µM lipid II and 1 mM MgCl_2_ in 20 mM Tris/HCl pH 7.5 with or without 150 mM NaCl as indicated for each experiment. Samples were incubated for 90 min at 37°C with shaking at 800 rpm. Reactions were stopped by boiling for 5 min. Digestion with cellosyl, reduction with sodium borohydride and analysis by HPLC were performed as described (Biboy *et al*., 2013).

#### FRET-based in vitro peptidoglycan synthesis assay in liposomes

For assays with liposomes, samples contained 20 mM Tris pH 7.5, 1 mM MgCl_2_ in a final volume of 50 µL. In this case, the same volume for each liposome preparation was added to the reactions, 10 µL, so that the total amount of labelled lipid II was present in every reaction. In these conditions, concentration of lipid II-Atto550 and lipid II-Atto647n would be 14.5 µM each, assuming no loss of lipids during sample preparation. The final concentration of enzymes, determined by densitometry of SDS-PAGE gels, were ∼0.59 µM for PBP1B^Ec^, ∼0.81 µM for PBP1B^Ab^, and ∼0.53 µM for PBP1B^Pa^. When indicated, activators LpoB(sol), LpoP^Ab^(sol), or LpoP^Pa^(sol) were added at a concentration of 2 µM. Reactions were started by the addition of lipid II at 12 µM and monitored by measuring fluorescence over a period of 60 min (or 90 min for PBP1B^Pa^ liposomes) at 37°C using a Clariostar plate reader (BMG Labtech, Germany), with emission measurements at 590 nm and 680 nm after excitation at 522 nm. When indicated ampicillin was added at 1 mM and moenomycin was added at 50 µM. Activity assays were performed immediately after preparation of liposomes was finished, as we noticed that some proteins could slowly start polymerization using the labelled lipid II. After reactions, samples were analysed by Tris-Tricine SDS-PAGE as indicated for detergents.

### Assays in supported lipid bilayers

#### Preparation of small unilamellar vesicles (SUVs) and proteoliposomes for SLB formation

Liposomes of *EcPL* lipids and proteoliposomes with reconstituted PBP1B^Ec^ were prepared as described previously by addition of beta-cyclodextrin to the solution of lipids and Triton X-100 detergent (DeGrip *et al*., 1998, Roder *et al*., 2011). Briefly, a thin lipid film of *E. coli* polar lipid extract was prepared by N_2_ assisted chloroform evaporation. After 2h of drying under vacuum the lipid film was re-hydrated to 5 mM (total phosphorus concentration) in 150 mM NaCl, 10 mM Tris-HCl, pH 7.4 supplemented with 20 mM Triton X-100. The suspension of lipids/detergent was extensively vortex, freeze/thawed for 5 cycles and sonicated using a water-bath sonicator for 10 min (on ice, to avoid lipids overheating upon sonication). To prepare proteoliposomes, full length PBP1B produced as described above and containing 0.05% Triton X-100 was mixed with a lipid-detergent suspension at the indicated ratio, usually 1:25000 (protein:lipids) and incubated for 10 min at RT. Incorporation of PBP1B^Ec^ into liposomes was achieved by addition of 2× excess of beta-cyclodextrin solution for 5 min (at RT) with subsequent 20-fold dilution in 20 mM Hepes, pH 7.4. The rapid depletion of detergent by addition of beta-cyclodextrin leads to formation of very small unilamellar vesicles with an average diameter of 18-25 nm and narrow size distribution (Roder *et al*., 2011).

To prepare liposomes with fluorescently labelled lipid II the extract of *E. coli* polar lipids was supplemented with 2 mol% solution of either lipid II-Atto550 or lipid II-Atto647n. The lipid film was treated similar as the film for preparation of proteoliposomes. Liposomes were also prepared by cyclodextrin-assisted extraction of Triton X-100.

#### Formation of polymer-supported lipid bilayers (SLBs) and reconstitution of PBP1B^Ec^ into a supported lipid membrane

To form polymer-supported lipid membranes the coverslips were functionalised beforehand with a dense PEG film, where the ends of the polymer brush were covalently modified with palmitic acid, which served as a linker to capture liposomes as described elsewhere (Roder *et al*., 2011). To perform a FRET assay on supported lipid membrane empty EcPL liposomes (i), liposomes with 2 mol% of either lipid II-Atto550 (ii) or lipid II-Atto647n (iii), and PBP1B^Ec^ proteoliposomes (iv) were mixed at equimolar ratio; and diluted by 20-fold with the 10 mM Tris pH 7.5 buffer directly in the reaction chamber. After 30 min of incubation at 37 °C the reaction chamber was washed 5 times by solution exchange. Proteoliposomes adsorbed on the surface were fused by the addition of 10% (w/v) PEG 8kDa solution (in water). The fusion reaction was carried for 15 min at 37°C, afterwards PEG solution was rigorously washed out. Fluidity and homogeneity of the lipid membrane were checked either with PE-Rhodamine dye (Avanti) or by addition of a His_6_-tagged (on the C-terminus) neutral peptide (CMSQAALNTRNSEEEVSSRRNNGTRHHHHHH) labelled with a single Alexa 488 fluorophore on its only Cys residue at the N-terminus to the EcPL membrane containing 0.1 mol% dioctadecylamine (DODA)-tris-Ni-NTA (Beutel *et al*., 2014).

#### FRET-based in vitro peptidoglycan synthesis assay in supported lipid bilayers using TIRF microscopy

Peptidoglycan synthesis reactions were carried out at 10 mM Tris pH 7.5 supplemented with 1 mM MgCl_2,_ with or without 1 mM Ampicillin and in the presence of 4 µM LpoB(sol). The reaction was started by addition of 4 µM of unlabelled lipid II. TIRF microscopy, using a set up described elsewhere(Baranova *et al*., 2020) was used to monitor an increase in FRET efficiency and spatial reorganization of FRET signal over the time course of PG synthesis. To detect real-time FRET on supported lipid membranes we used the so-called “acceptor photobleaching approach” where a region of interest of about 10×10 µm was photobleached in the acceptor channel (lipid II-Atto647n) and the increase in fluorescence intensity of the donor (lipid II-Atto550) was recorded within a delay of 1s. The FRET efficiency was calculated as described(Loose *et al*., 2011). Briefly, donor intensity levels were calculated before (I^D^) and after photobleaching (I^D,pb^) using intensity measurements in ImageJ. FRET efficiency was calculated using Equation 1:

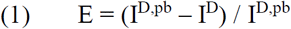

For time-course measurements (Figure 4D) the acceptor signal (lipid II-Atto647n) was photobleached every minute after the initiation of the reaction (the data point at time 0 corresponds to addition of unlabelled lipid II). For each time point a new region of interest in the same chamber was photobleached, and the change in the donor intensity was recorded to calculate FRET efficiency using Eq.1.

To have a control on the lipid membrane integrity during PG synthesis the phospholipid DODA-tris-Ni-NTA (Beutel *et al*., 2014) was included during reconstitution at a 0.1 mol% ratio. DODA-tris-Ni-NTA was then visualized using a His_6_-containing peptide (CMSQAALNTRNSEEEVSSRRNNGTRHHHHHH) labelled with Alexa488 on its single Cys residue, which we added in the same experiment in which we performed FRET analysis. To compare the fluidity and the immobile fraction of lipid II-Atto647n before and after 1 h of the synthesis reaction with the fluidity of phospholipids in the lipid membrane, the same region of interest was photobleached with a laser first at 640 nm and afterwards at 480 nm.

#### In vitro peptidoglycan synthesis assay using radiolabelled lipid II on supported lipid bilayers

To assay PG synthesis on supported lipid bilayers (SLBs) using radioactively labelled lipid II, we first reconstituted PBP1B^Ec^ on SLBs containing *E. coli* polar lipid extract and a 1:10^5^ PBP1B^Ec^ to lipid molar ratio, as described above. Due to the low density of the enzyme, several 1.1 cm^2^ chambers were assayed for every condition in order to accumulate a measurable signal. In every chamber, reactions were started by addition of 10 µM [^14^C]-labelled lipid II and 4 µM LpoB(sol) in a total volume of 100 µL per chamber. The synthesis reaction was carried out in 10 mM Tris pH 7.5, 1 mM MgCl_2_. The chambers were incubated overnight (∼16h) at 37°C, covered with parafilm. Reactions were stopped by addition of 100 µM moenomycin. To digest the produced peptidoglycan, cellosyl was added at 0.05 g/L, in the presence of 0.3% triton X-100. After 1h incubation at 37°C, samples from 6 Chambers were pooled in an Eppendorf tube, concentrated using a speed-vac evaporator, reduced using sodium borohydride and analysed by HPLC as described above. For the experiment to determine lipid II incorporation and the localisation of the produced PG, before addition of moenomycin, chambers were washed by removal of 50 µL of buffer and addition of 50 µL of fresh buffer while mixing. This was repeated 5 times. The removed volume from each wash was pooled and treated the same as the samples left in the chamber.

#### Single molecule tracking and analysis

To perform single molecular tracking, MGC-^64^PBP1B-his C77S C795S was labelled with the photostable far-red dye Dy647N as described above and then reconstituted into a polymer-supported lipid membrane as described elsewhere (Roder *et al*., 2011, Roder *et al*., 2014). Single molecule tracking experiments were performed at a low protein to lipid molar ratio (1:10^−6^). At this ratio, supported lipid membrane was largely homogeneous with the lowest immobile fraction from all the ratios tested (Figure 3 – figure supplement 1). The single-molecule motion of PBP1B was measured prior and after the addition of 1.5 µM lipid II after 15 min ex situ incubation, in the presence of 10 mM Hepes pH 7.4, 150mM NaCl, 1 mM MgCl_2_ buffer and in the absence of LpoB(sol). The localization and tracking of PBP1B particles was performed by the SLIMfast software (Serge *et al*., 2008). To ensure that non-specifically stuck PBP1B particles did not contribute to the measured diffusion coefficient the localized movies, the immobile particles were excluded using the DBSCAN spatial clustering algorithm (Sander *et al*., 1998) with the following clustering parameters: a search area of 100 nm, the minimal time window of 30 frames at 65 ms/frame acquisition time. The displacement distributions for active PBP1B (in the presence of lipid II) was compared to the displacement distribution of PBP1B before lipid II addition by fitting the two-component Rayleigh distribution and comparing the weighted contribution of each population. The mean-squared displacement was fitted to each individual trajectory longer than 650 ms (10 frames). Each MSD curve was fitted with a linear fit considering max 30% of the lag-time for each trajectory.

#### FRAP analysis

To control membrane fluidity upon the reconstitution of the transmembrane PBP1B (Figure 3 – figure supplement 1 and Figure 4 – figure supplement 1) and fluidity of lipid II Atto-647n during peptidoglycan synthesis (Figure 4E-F) we used a Matlab-based GUI frap_analysis (Jönsson, 2020) in details described elsewhere (Jönsson *et al*., 2008). This code allows to quantify the contribution of the immobile fraction to the estimated diffusion coefficient, and particularly suitable for the analysis of 2D diffusion with the photobleaching contribution during the recovery.

## ACKNOWLEDGEMENTS

We thank Alexander Egan (Newcastle University) for purified proteins LpoB(sol) and LpoP^Pa^(sol), Oliver Birkholz and Jacob Piehler for their help with PBP1B reconstitution into polymer-SLBs and initial guidance on single particle tracking. We also acknowledge Changjiang You for providing tris-DODA-NTA reagent. This work was funded by the BBSRC grant BB/R017409/1 (to W.V.), by the European Research Council through grant ERC-2015-StG-679239 (to M.L.), and long-term fellowships HFSP LT 000824/2016-L4 and EMBO ALTF 1163-2015 (to N.B.).

## FIGURE LEGENDS

**Figure 1 - figure supplement 1.**
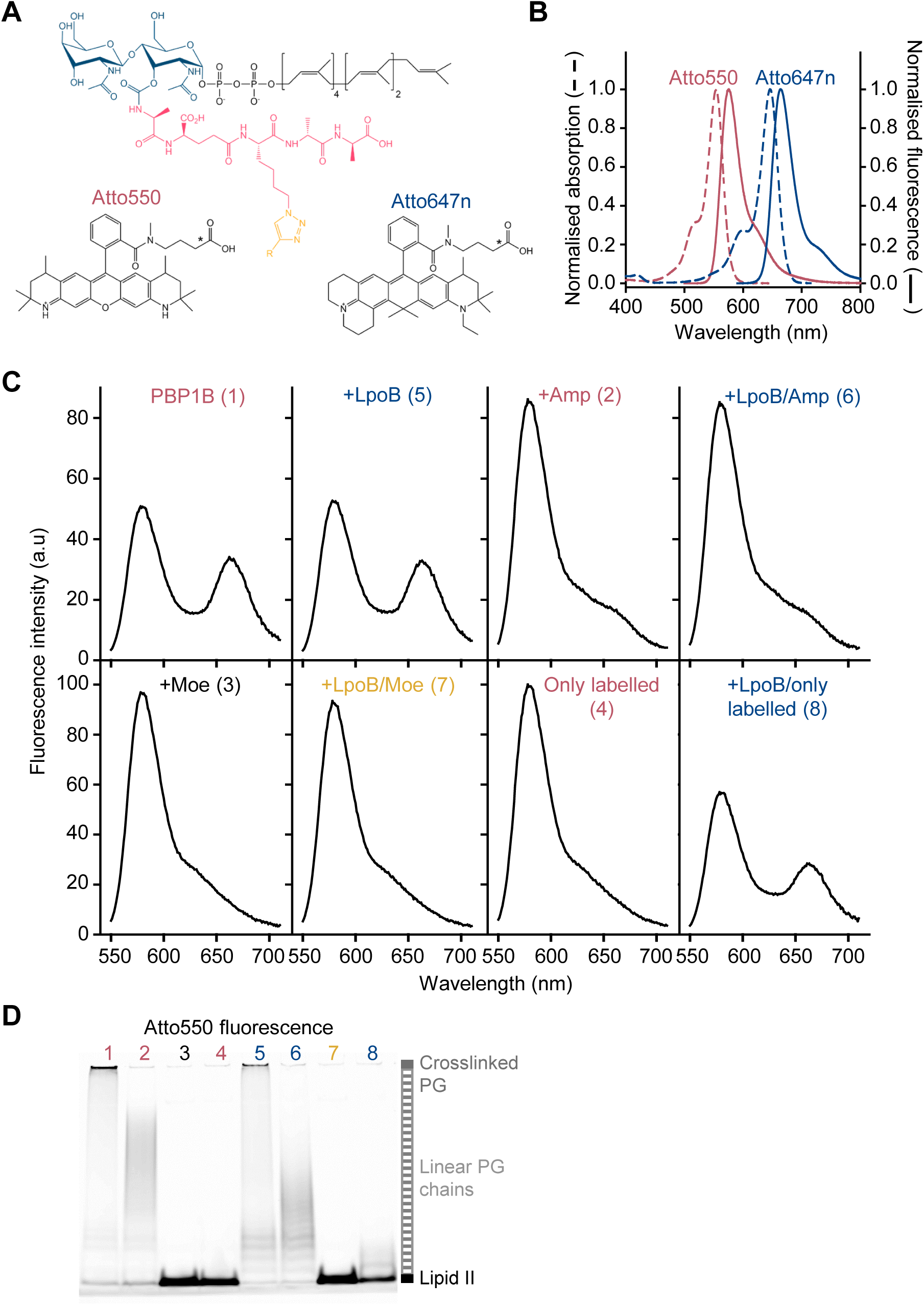
FRET assay to monitor PG synthesis in real time. (**A**) Chemical structures of lipid II analogues used for the FRET assay. R corresponds to Atto550n (donor) or Atto647n (acceptor) in the corresponding analogue. The chemical structures of alkyne versions of Atto550 and Atto647n probes that were used for derivatization are not published. Therefore the carboxylic variants are depicted here with an asterisk indicating where the alkyne versions diverge. (**B**) Absorbance (dashed lines) and fluorescence emission (solid lines) spectra for Atto550 (red lines) and Atto647n (blue lines). (**C**) Fluorescence emission spectra taken at the end (t=1 h) of the reactions of *E. coli* PBP1B shown in Figure 1B (t=60 min). (**D**) The same gel depicted in Figure 1D, but scanned using the donor fluorescence (Atto550n).

**Figure 2 - figure supplement 1.**
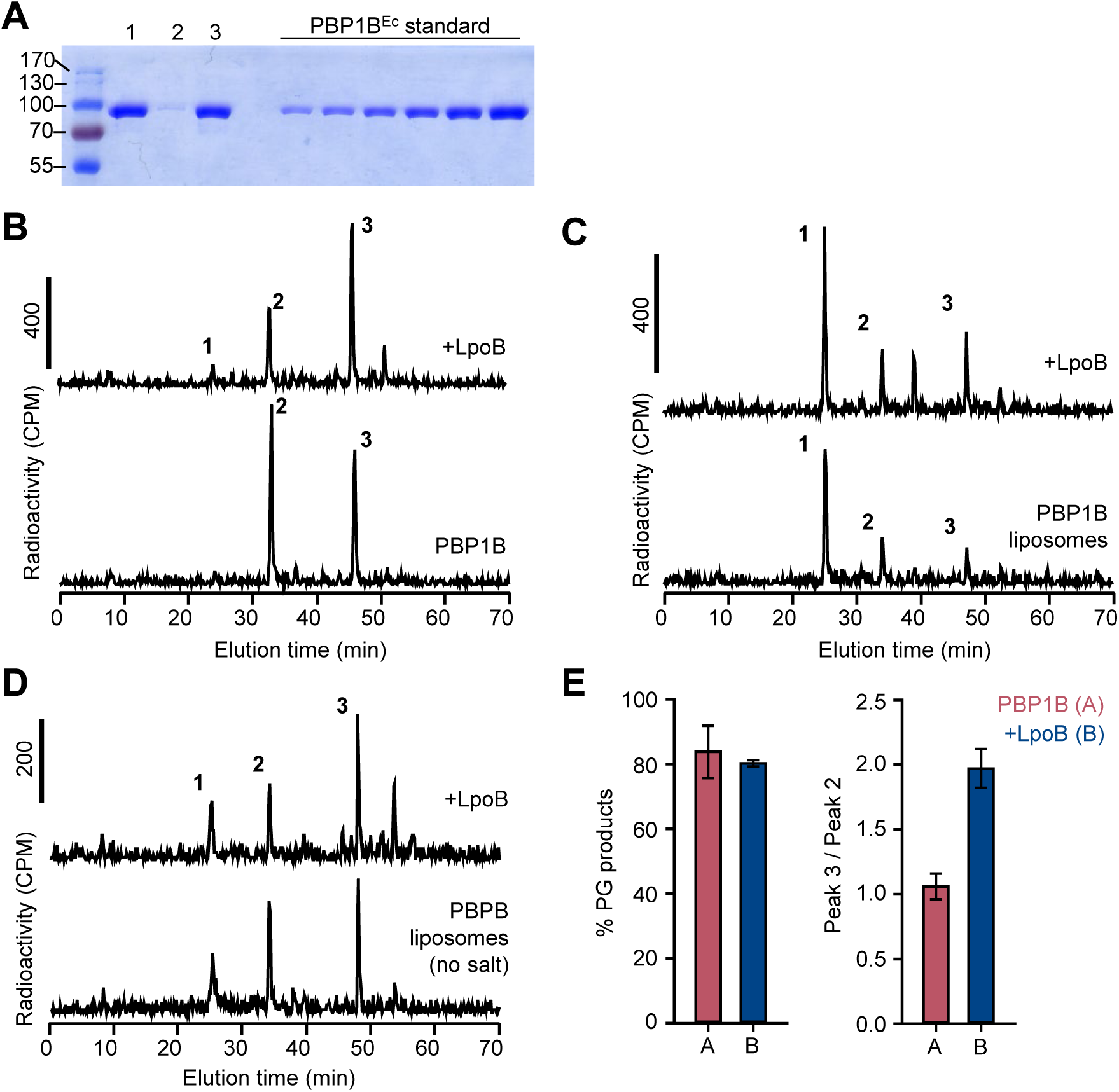
Activity of membrane-reconstituted PBP1BEc is optimal in *E. coli* polar lipids at low ionic strength. (**A**) Representative SDS-PAGE analysis of the reconstitution of PBP1B^Ec^ in liposomes made of *E. coli* polar lipids at a 1:3000 mol:mol protein:lipid ratio. After reconstitution, proteoliposome samples (lane 1) were centrifuged at low speed to remove aggregates and both pellet and supernatant samples were analysed (lanes 2 and 3, respectively). The supernatant was subsequently used for PG synthesis reactions. A gradient of PBP1B^Ec^ (0.25, 0.41, 0.62, 0.82, 1.23 and 1.65 µg) was loaded as a standard to estimate protein concentration by densitometry. (**B**), (**C**) and (**D**) Representative chromatograms showing the muropeptide analysis of PG produced by detergent-solubilised PBP1B^Ec^ (**B**) or liposome-reconstituted PBP1B^Ec^ in the presence or absence of NaCl (**C** and **D**, respectively). The concentration of PBP1B^EC^ was 0.5 µM and, if added, that of LpoB(sol) was 2 µM LpoB(sol). The reaction buffer contained 150 mM NaCl in **B** and **C**. Samples were incubated at 37 °C for 60 min in **B** and 90 min in **C** and **D**. The labelled peaks correspond to the muropeptides shown in Figure 1E. (**E**) Quantification of the total amount of radioactivity incorporated into PG (left) or the ratio between the radioactivity of peaks 3 and 2 (indicative of the degree of crosslinking of the PG, right) for activity assays for PBP1B^Ec^ in liposomes in the same conditions as in **D**. Values are mean ± SD (or variation) of at least two reactions.

**Figure 2 - figure supplement 2.**
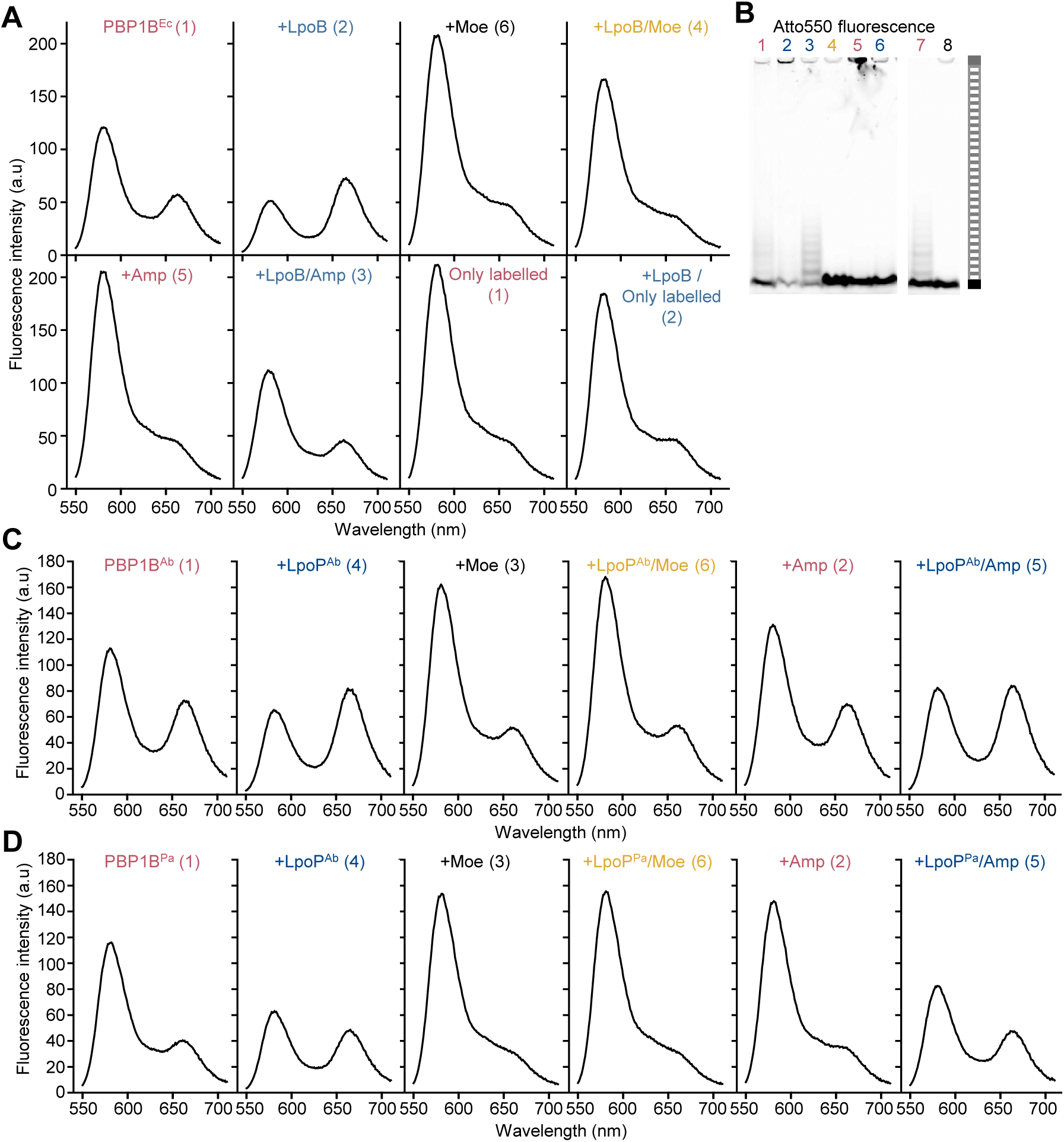
The FRET assay for PG synthesis can be adapted for reactions on liposomes. **(A)** Spectra corresponding to *E. coli* PBP1B reactions shown in Figure 2C, taken at t=60 min. (**B**) The same gels depicted in Figure 2C, but scanned using the donor fluorescence (Atto550n). (**C**) Spectra corresponding to *A. baumannii* PBP1B reactions shown in Figure 2E, taken at t=60 min. (**D**) Spectra corresponding to *P. aeruginosa* PBP1B reactions shown in Figure 2G, taken at t=90 min.

**Figure 2 - figure supplement 3.**
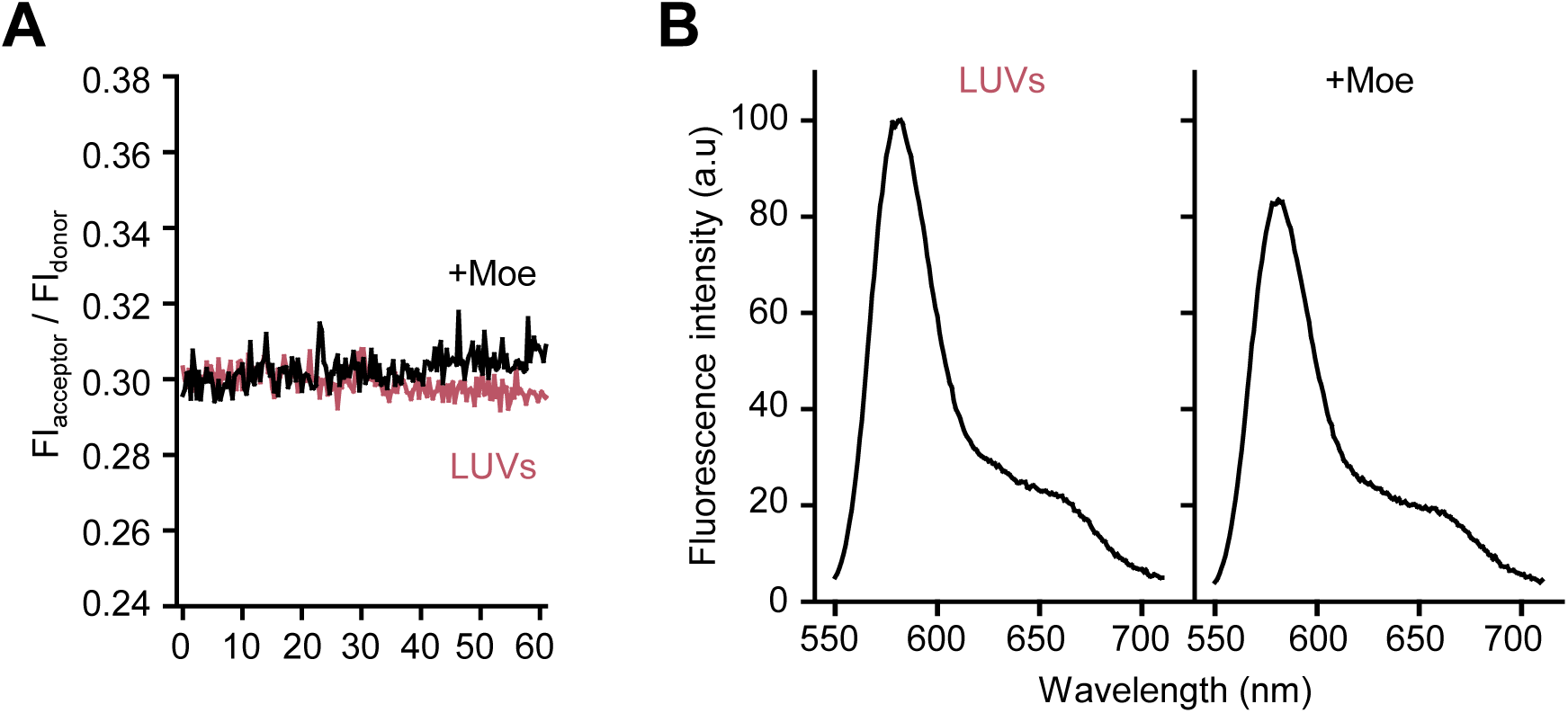
Moenomycin does not affect FRET on liposomes with lipid II-Atto550 and lipid II-647 in the absence class A PBPs. (**A**) EcPL liposomes incorporating an equimolar amount of lipid II-Atto550 and lipid II-Atto647n at 0.5%mol of the total lipid contents where incubated in the presence of 12 µM lipid II and in the presence (black line) or absence (red line) of 50 µM moenomycin for 60 min at 37 °C while monitoring FRET as indicated in materials and methods. (**B**) Fluorescence spectra for the samples described in **A** at the end of the incubation period (t=60 min).

**Figure 2 - figure supplement 4.**
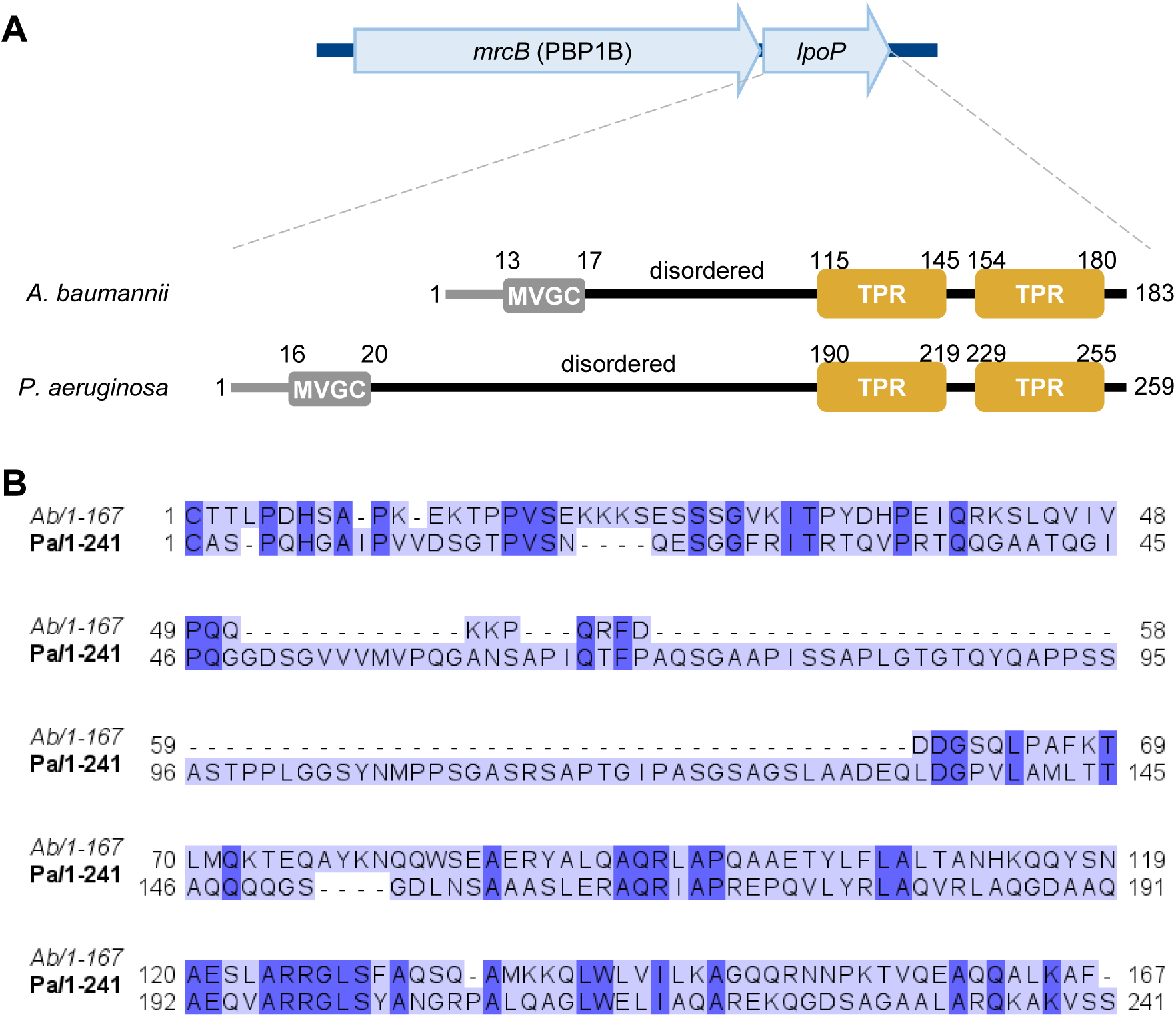
Amino acid sequence comparison between LpoP homologues from *A. baumannii* and *P. aeruginosa*. (**A**) In the genomes of *A. baumanni* and *P. aeruginosa* the gene encoding LpoP is present within the same operon as the gene encoding their cognate PBP1B. Both LpoP proteins are predicted lipoproteins with a disordered region between the N-terminal Cys and the C-terminal globular domain containing the tetratricopeptide repeats (TPR). LpoP^Ab^ has a shorter disordered linker than LpoP^Pa^. (**B**) Sequence comparison between the globular regions of LpoP^Ab^ (Ab) and LpoP^Pa^ (Pa). Proteins sequences (minus the signal peptides) were aligned using T-COFFEE EXPRESSO and the resulting alignment was visualized using JALVIEW. Residues conserved in both proteins are highlighted in a darker colour.

**Figure 2 - figure supplement 5.**
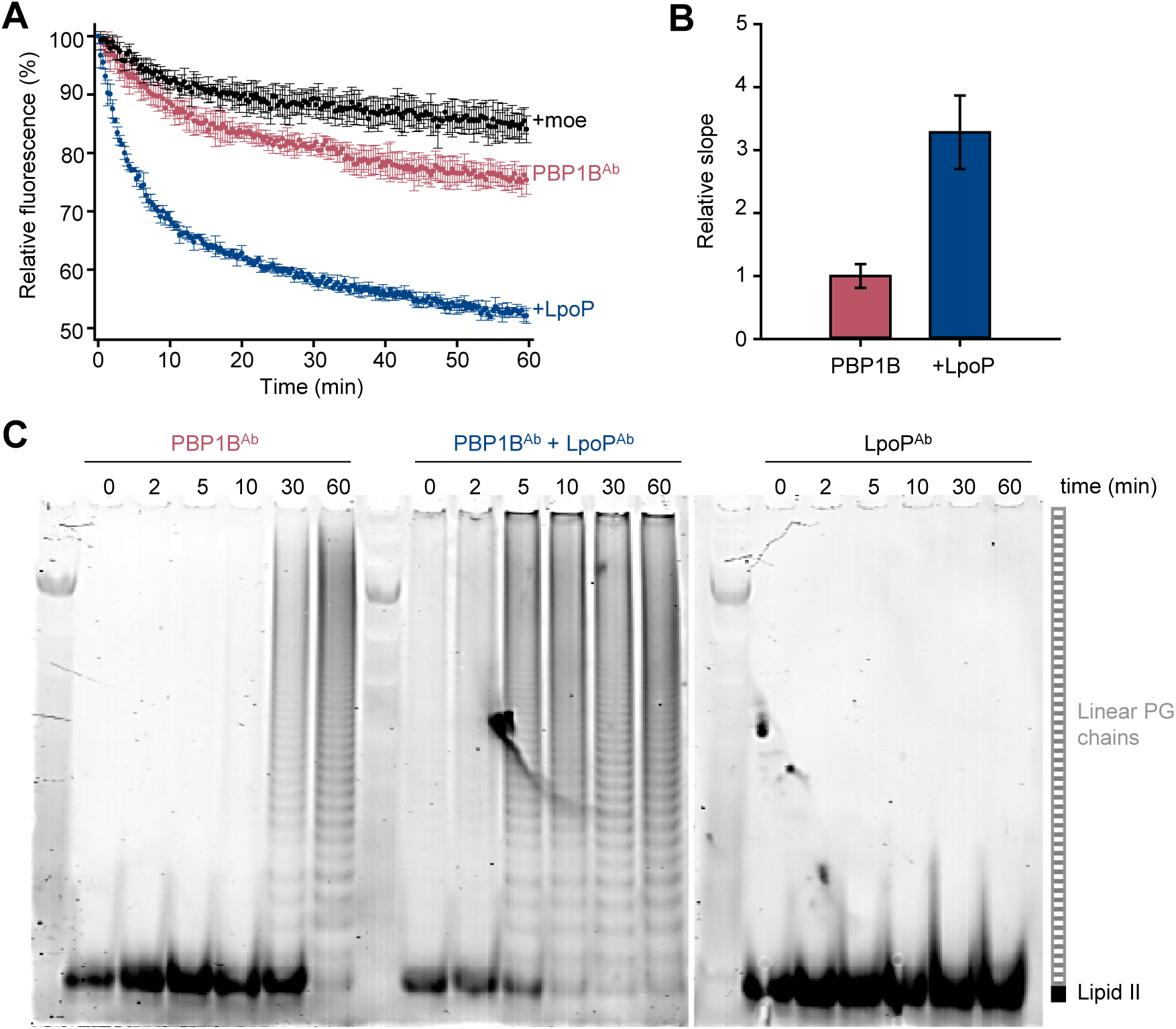
LpoP^Ab^ stimulates the glycosyltransferase activity of PBP1BAb. (**A**) Real-time glycosyltransferase activity assays using dansyl-lipid II and detergent-solubilised *A. baumannii* PBP1B (PBP1B^Ab^). PBP1B^Ab^ (0.5 µM) was mixed with 10 µM dansyl-lipid II in the presence or absence of soluble 0.5 µM *A. baumannii* LpoP (LpoP^Ab^(sol)). A control was performed by adding 50 µM moenomycin (black). Each data point represeants mean ± SD of 3 independent experiments. (**B**) Averaged initial slopes from reaction curves in **A**. Values are normalised relative to the slope in the absence of activator and are mean ± SD of 3 independent experiments. (**C**) Time-course GTase assay by SDS-PAGE followed by fluorescence detection. Detergent-solubilised PBP1B^Ab^ was incubated with 5 µM lipid II-Atto550 and 25 µM unlabelled lipid II in the presence or absence of 1.5 µM LpoP^Ab^(sol). Reactions contained 1 mM Ampicillin to block transpeptidation. Aliquots were taken at the indicated times (in min), boiled and analysed by SDS-PAGE. A control in which only LpoP^Ab^(sol) was present is also shown.

**Figure 2 - figure supplement 6.**
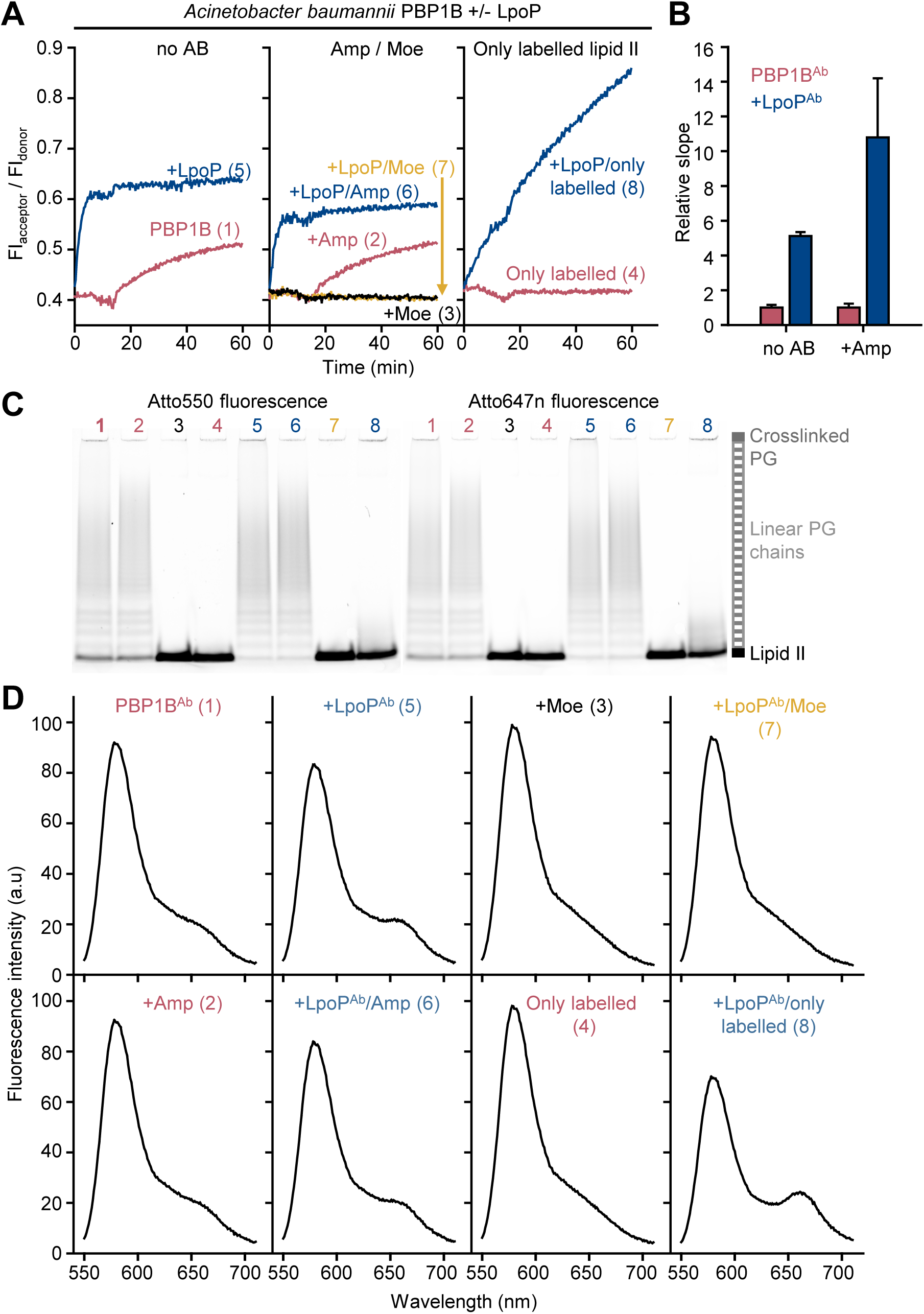
PG synthesis activity of *A. baumannii* PBP1B in the presence of Triton X-100 followed by FRET. (**A**) Representative FRET curves for activity assays using detergent-solubilised *A. baumannii* PBP1B (PBP1B^Ab^). PBP1B^Ab^ (0.5 µM) was mixed with unlabelled lipid II, Atto550-labelled lipid II and Atto647n-labelled lipid II at a 1:1:1 molar ratio (5 µM of each), in the presence or absence of 2 µM soluble *A. baumannii* LpoP (LpoP^Ab^(sol)). Controls were performed by adding 50 µM moenomycin in the absence (black) or presence (yellow) of LpoP^Ab^(sol). Reactions were performed without antibiotic (left), with 1 mM ampicillin (middle), or in the absence of unlabelled lipid II (right). The numbers indicate the corresponding lane of the gel in **C**. Samples were incubated for 60 min at 30°C. (**B**) Averaged initial slopes from reaction curves obtained by the FRET assay for detergent-solubilised PBP1B^Ab^, in the presence (blue) or absence (red) of LpoP, and in the presence or absence of ampicillin. Values are normalised relative to the slope in the absence of activator for each condition and are mean ± SD of 2 independent experiments. (**C**) Aliquots after reactions in **A** were boiled and analysed by SDS-PAGE followed by fluorescence detection. (**D**) Fluorescence emission **s**pectra taken after reactions in **A** (t=60 min).

**Figure 2 - figure supplement 7.**
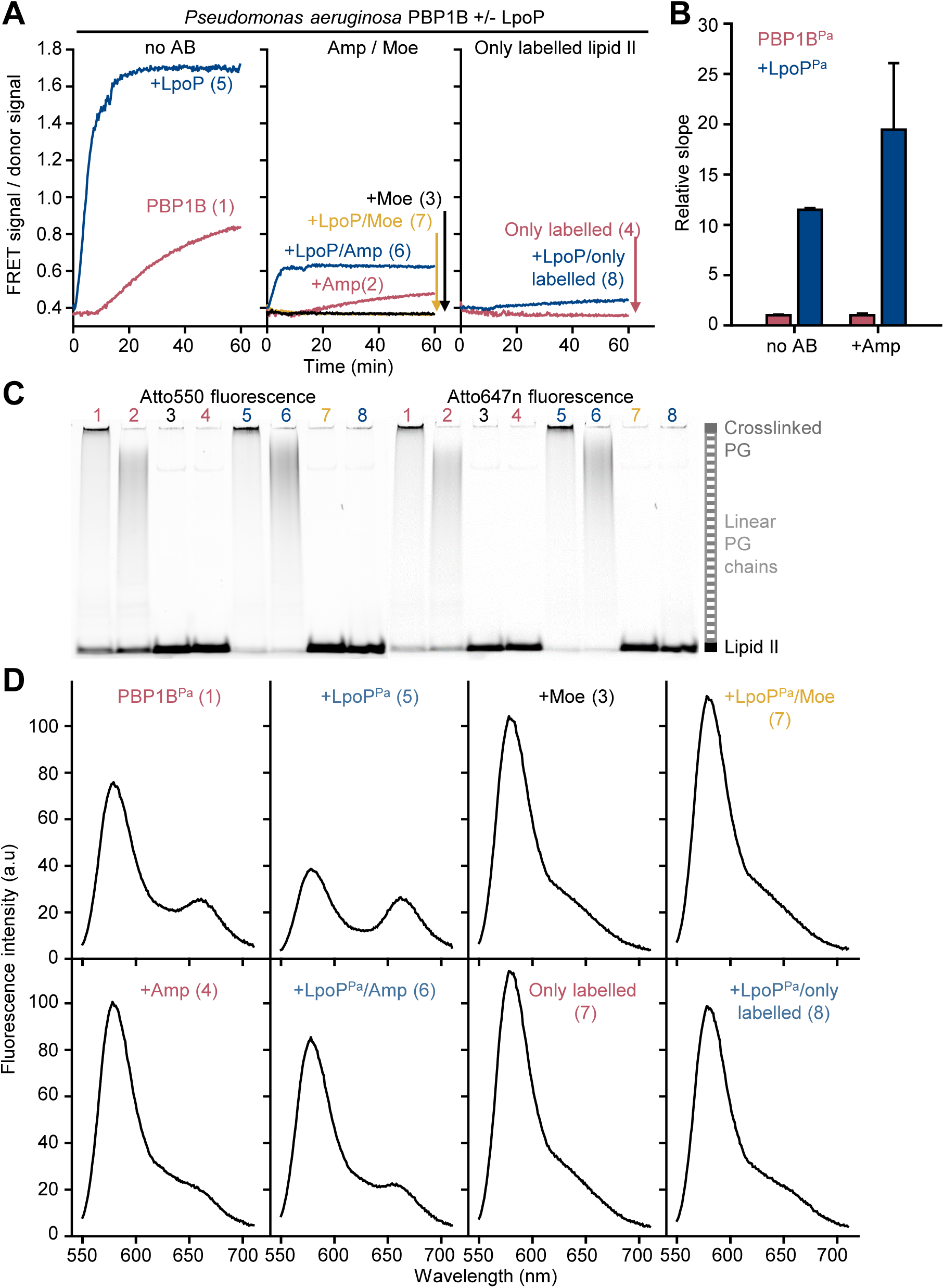
PG synthesis activity of *P. aeruginosa* PBP1B in the presence of Triton X-100 followed by FRET. (**A**) Representative FRET curves for activity assays using detergent-solubilised *P. aeruginosa* PBP1B (PBP1B^Pa^). PBP1B^Pa^ (0.5 µM) was mixed with unlabelled lipid II, Atto550-labelled lipid II and Atto647n-labelled lipid II at a 1:1:1 molar ratio (5 µM of each), in the presence or absence of 2 µM soluble *P. aeruginosa* LpoP (LpoP^Pa^ (sol)). Controls were performed by adding 50 µM moenomycin in the absence (black) or presence (yellow) of LpoP^Pa^(sol). Reactions were performed without of antibiotic (left panel), with 1 mM ampicillin (middle panel), or in the absence of unlabelled lipid II (right panel). The numbers indicate the corresponding lane of the gel in **C**. Samples were incubated for 90 min at 37°C. (**B**) Averaged initial slopes from reaction curves obtained by the FRET assay for detergent-solubilised PBP1B^Pa^, in the presence (blue) or absence (red) of LpoP, and in the presence or absence of ampicillin. Values are normalised relative to the slope in the absence of activator for each condition and are mean ± SD of 2-3 independent experiments. (**C**) Aliquots after reactions in **A** were boiled and analysed by SDS-PAGE followed by fluorescence detection. (**D**) Fluorescence emission **s**pectra taken after reactions in **A** (t=90 min).

**Figure 3 - figure supplement 1.**
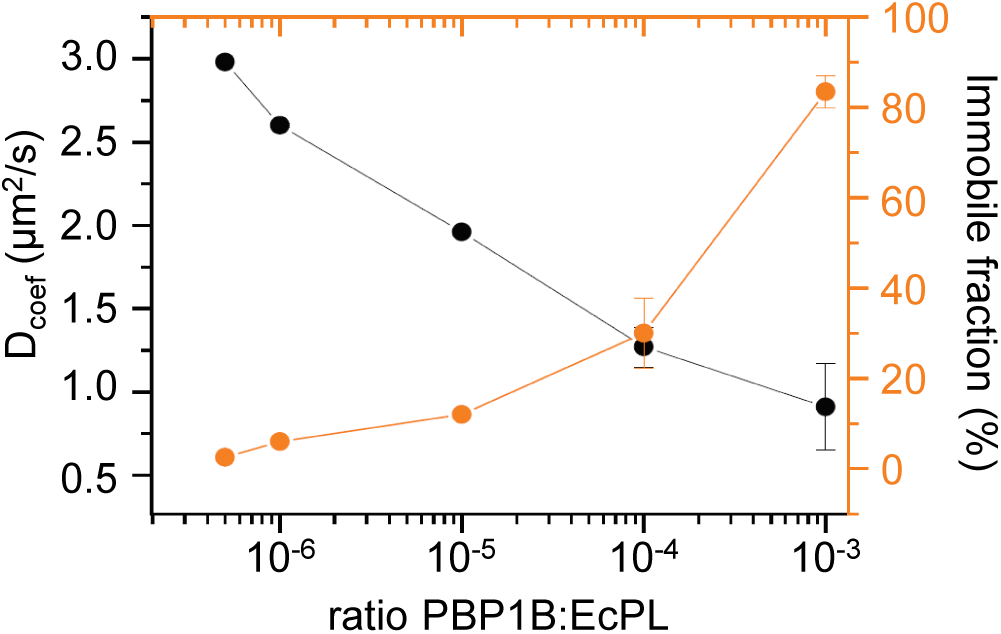
Control of membrane fluidity and integrity upon reconstitution of *E. coli* PBP1B. (**A**) The fluidity of supported lipid bilayers is reduced when increasing PBP1B^Ec^ density. The diffusion of phospholipid probe DOPE-rhodamine in the polymer-supported SLB was monitored by FRAP at different densities of PBP1B. The fluidity of the membrane decreased (black line) while the immobile fraction increased (orange line) with higher protein densities.

**Figure 3 - figure supplement 2.**
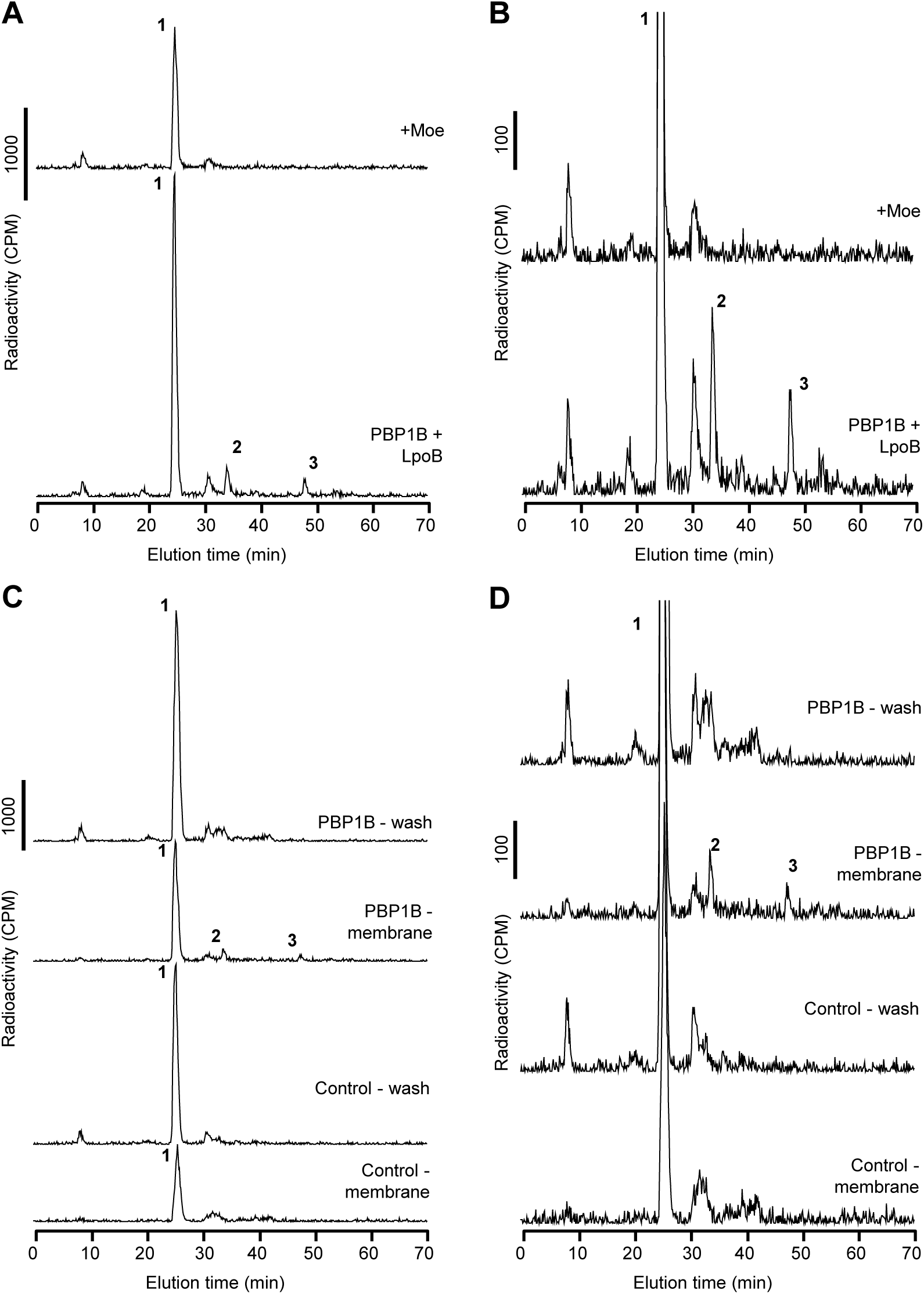
*E. coli* PBP1B is active after reconstitution in supported lipid bilayers. (**A**) and (**B**) PBP1B^Ec^ was reconstituted on supported lipid bilayers prepared with *E. coli* polar lipid extract in 1.1 cm^2^ chambers. The protein to lipid ratio was 1:10^5^ (mol:mol). Reactions were started by adding 1 nmol of radiolabelled lipid II per chamber, in the presence of LpoB(sol) (4 µM) moenomycin (100 µM). Three chambers were prepared for each condition and samples were combined before the analysis. Chambers were incubated overnight at 37 °C and the reaction was stopped by adding moenomycin. Cellosyl and Triton X-100 were added to solubilize the membranes and digest the PG product. The resulting muropeptide samples were concentrated, reduced with sodium borohydride and analysed by HPLC. Full chromatograms are shown in **A**, while zoomed-in chromatograms are shown in **B**. (**C**) and (**D**) PG synthesis occurs only in the membrane fraction of SLBs. PBP1B^Ec^ was reconstituted on SLBs as in **A** and **B**. In addition, control chambers were prepared without PBP1B. Chambers were incubated over night to allow for PG synthesis and then washed with fresh buffer. The washes and chambers (membranes) were treated and analysed as described for **A** and **B**. Five chambers were combined for reactions with PBP1B^Ec^, and four chambers for control reactions. Full chromatograms are shown in **C**, while zoomed-in chromatograms are shown in **D**. The labelled peaks in all chromatograms correspond to the muropeptides shown in Figure 1F.

**Figure 4 - figure supplement 1.**
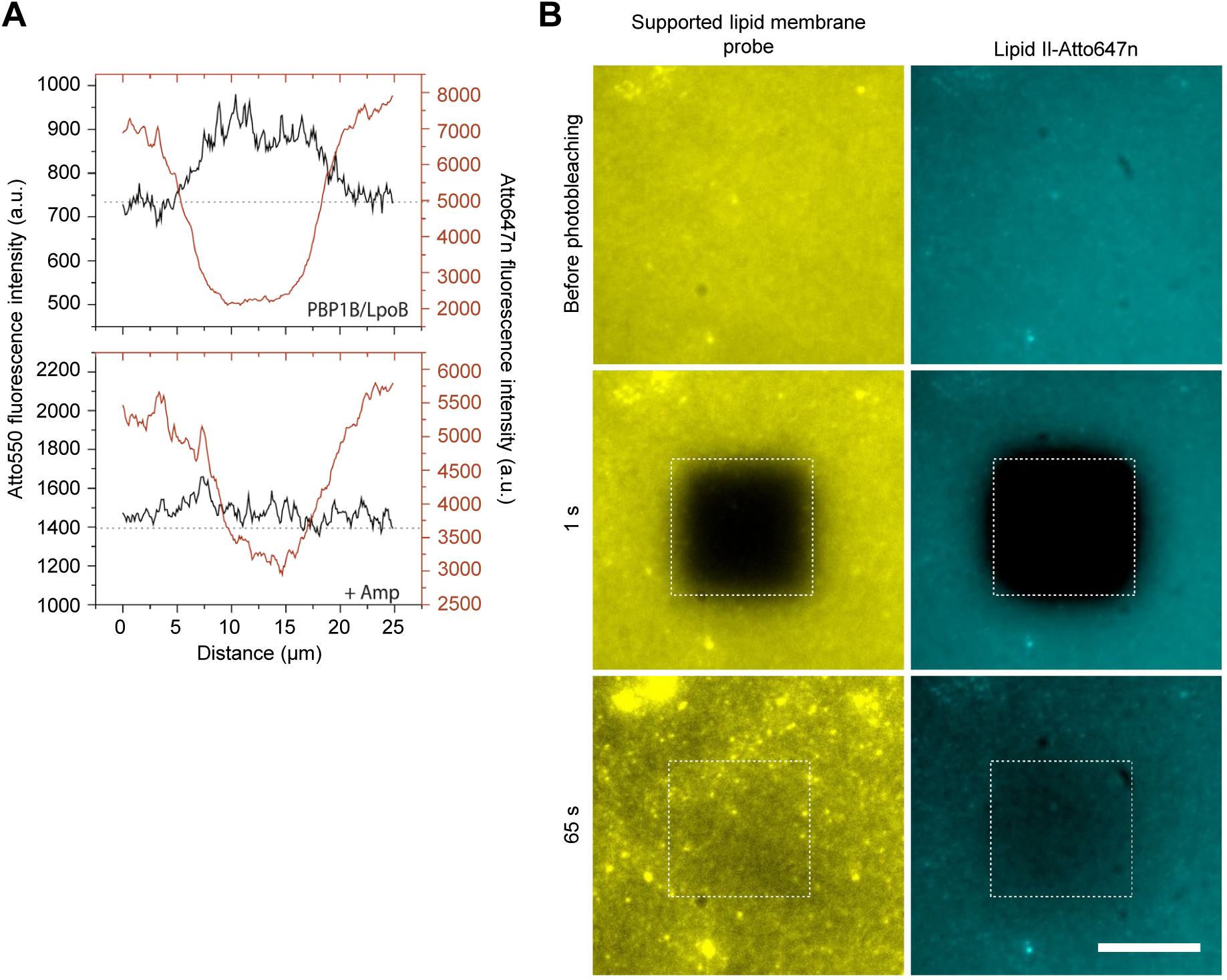
Control of membrane fluidity and integrity during the FRET assay. (**A**) Fluorescence intensity profiles 1s after photobleaching taken from the images depicted on Figure 4B. (**B**) Montage comparing the recovery of fluorescence after photobleaching of a tracer (DODA-tris-Ni-NTA plus a His_6_-tagged peptide labelled with AlexaFluor 488) with the one of lipid II-Atto647n on a supported lipid bilayer containing PBP1B at a 1:10^5^ protein:lipid (mol:mol) ratio. The assay was performed after a PG synthesis reaction carried out for 1.5 h. The fact that fluorescence is recovered for both, indicates that the membrane remains fluid while lipid II stays diffusive after the synthesis reaction.

**Movie 1. Single-molecule imaging of PBP1B on supported lipid bilayers**. PBP1B^Ec^- Dy647 was reconstituted in EcPL SLBs at a 1:10^6^ (mol:mol) protein to lipid ratio and was tracked using single-molecule TIRF before or after the addition of 1.5 µM lipid II. Images were taken with a rate of 62 ms per frame.

**Movie 2. FRET assay on supported lipid bilayers**. PBP1B^Ec^ was reconstituted in EcPL SLBs at a 1:10^5^ (mol:mol) protein to lipid ratio along lipid II-Atto647, lipid II-Atto550. Membranes were incubated with 5 µM lipid II in the presence or absence of 1 mM ampicillin. To detect FRET, the fluorescence of the acceptor Atto647n was bleached within a region. In the subsequent frame the fluorescence of Atto550 increased indicating the presence of FRET. In the presence of ampicillin this increase did not happen.

**Supplementary File 1:** table of oligonucleotides used in this study.

**Supplementary Table 1.**
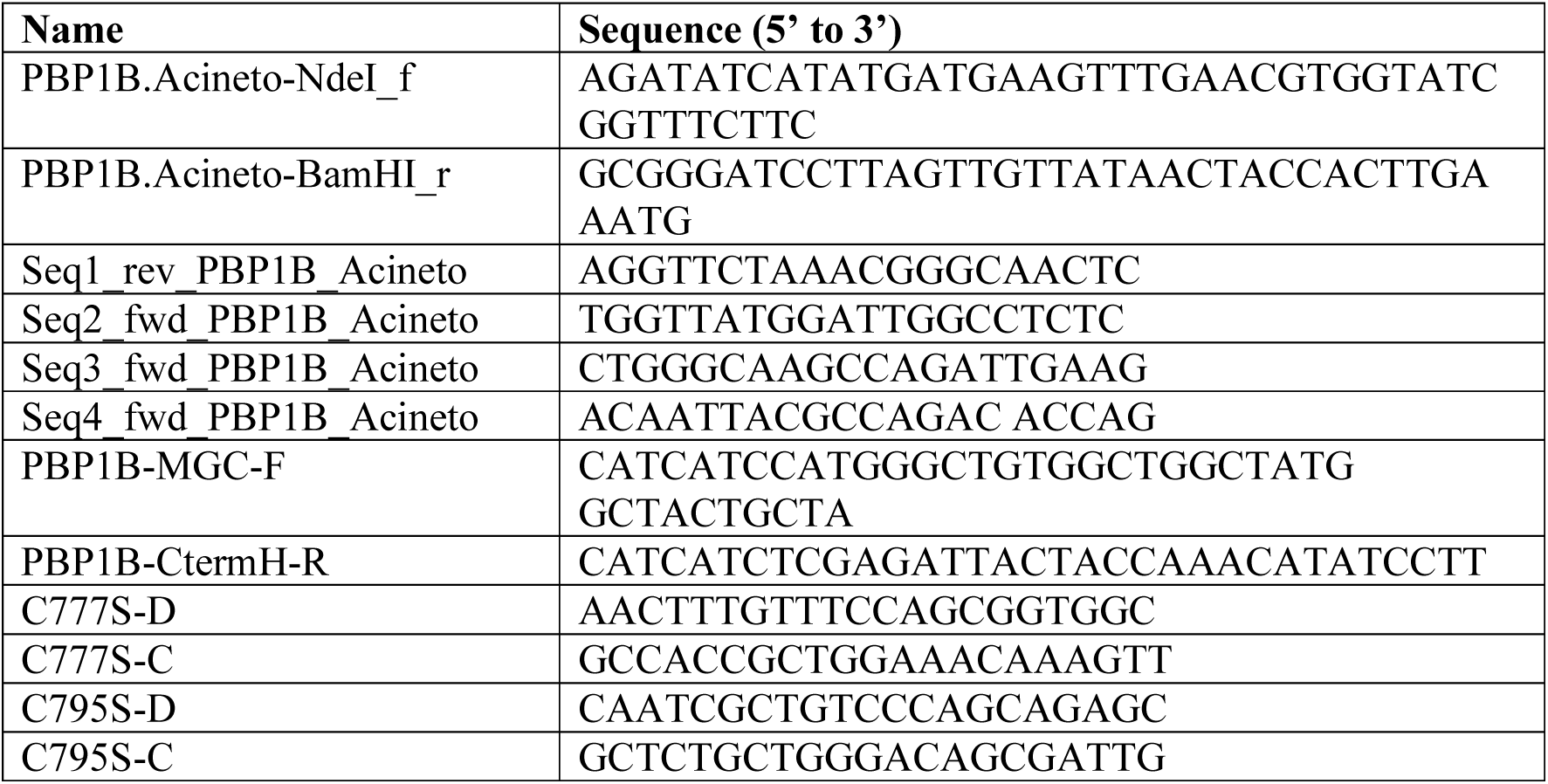
Oligonucleotides used in this work.

